# flowEMMi: An automated model-based clustering tool for microbial cytometric data

**DOI:** 10.1101/667691

**Authors:** Joachim Ludwig, Christian Höner zu Siederdissen, Zishu Liu, Peter F Stadler, Susann Müller

**Affiliations:** Department of Environmental Microbiology, Research Group Flow Cytometry, Helmholtz Centre for Environmental Research, Permoserstraße 15, 04318 Leipzig, Germany; Department of Computer Science, University Leipzig, Härtelstr. 16-18, 04107 Leipzig, Germany

**Keywords:** Flow cytometry, Clustering, Data analysis, Statistical analysis, Microbial communities, Expectation-Maximization

## Abstract

**Background:** Flow cytometry (FCM) is a powerful single-cell based measurement method to ascertain multidimensional optical properties of millions of cells. FCM is widely used in medical diagnostics and health research. There is also a broad range of applications in the analysis of complex microbial communities. The main concern in microbial community analyses is to track the dynamics of microbial subcommunities. So far, this can be achieved with the help of time-consuming manual clustering procedures that require extensive user-dependent input. In addition, several tools have recently been developed by using different approaches which, however, focus mainly on the clustering of medical FCM data or of microbial samples with a well-known background, while much less work has been done on high-throughput, online algorithms for two-channel FCM.

**Results:** We bridge this gap with `flowEMMi`, a model-based clustering tool based on multivariate Gaussian mixture models with subsampling and foreground/background separation. These extensions provide a fast and accurate identification of cell clusters in FCM data, in particular for microbial community FCM data that are often affected by irrelevant information like technical noise, beads or cell debris. `flowEMMi` outperforms other available tools with regard to running time and information content of the clustering results and provides near-online results and optional heuristics to reduce the running-time further.

**Conclusions:** `flowEMMi` is a useful tool for the automated cluster analysis of microbial FCM data. It overcomes the user-dependent and time-consuming manual clustering procedure and provides consistent results with ancillary information and statistical proof.

## Background

Flow cytometry (FCM) is a high-throughput technology to measure multidimensional optical properties of single cells. Flow cytometry is widely used in medical diagnosis and health research but there is also a large area of applications in the context of complex microbial communities. Microbial communities are present everywhere in our environment. They are also used in biotechnological applications e.g. for the treatment of waste water, the production of biogas or the manufacturing of platform chemicals. Here, FCM can be used for process monitoring such as for testing drinking water quality, process control and process improvement [1, 2, 3, 4, 5]. Natural systems can also be well described by flow cytometry and ecological measures such as diversity and stability indices that were recently established [6, 7, 8]. Flow cytometry was also already used to analyze the mice gut microbiome [9] and the human oral microbiome [10, 11].

The main concern in all of these applications is to follow microbial population [12] or microbial community structure variations. Even machine learning methods have been tested to identify exclusive strains in cytometrically measured in-silico communities [13]. As the generation times of microbial cells are very short and change population and community structures rapidly and thus also their interactions with environmental surroundings, information about structure variations need to be obtained in a very short time and in an automated way. Bioinformatics tools such as `flowCHIC` [14, 15] and `flowCyBar` [16, 17] were developed to reveal insights into microbial community variations.

While `flowCHIC` is an automated approach based on whole dot plot pixel densities and can be used to reveal pairwise structural variations between microbial communities, `flowCyBar` is based on gate/cell cluster information and provides insight into community structures based on numbers of subcommunities, the position of subcommunities within the dot plot and the number of cells inside subcommunities. `flowCyBar` allows to follow community evolution and if environmental parameters are involved in the evaluation pipeline, correlation analyses between those and subcommunity cell numbers can be performed in order to reveal functional dependencies. Subcommunities of interest can be flow sorted which allows further cell analysis employing next generation sequencing or proteomic approaches. Therefore, `flowCyBar` is an essential tool to determine cytometric community characteristics.

To perform the `flowCyBar` analysis, subcommunities have to be clustered according to their optical properties. These subcommunities are likely to have a certain function within biological processes and show correlations to certain environmental (abiotic) factors that can be revealed by using `flowCyBar`. The clustering of these subcommunities is the only step in the evaluation pipeline which is still performed manually in an experience-based and time-consuming way due to the high complexity of the data. Different from standard cytometric data of human samples, where cells are usually differentiated using a variation of labeled antibodies, and of different fluorescent excitations and emissions (resulting in only two or three different subpopulations per each 2D-plot) in bacterial flow cytometry the number of subcommunities can increase to up to 30 in each 2D-plot. Only two parameters (usually a nucleic acid dye and FSC) are sufficient to resolve bacterial community structures and follow their dynamics. The appearance of dozens of different clusters within only two dimensions is only known for bacterial samples and requires specialized evaluation procedures. These clusters provide information on cell abundance changes and anticipated cells can further be processed after cell sorting.

The automatic definition of that many gates in a 2D-plot is a bottleneck that cannot be solved by existing tools with satisfactory precision. To alleviate this issue, we developed a statistical model-based approach with as few as possible parameters (that *require* user control) that fulfills all the requirements on the outcomes of the clustering procedure of microbial community data.

Therefore, the approach (i) regards only two channels, (ii) recognizes typically between 10 to 20 clusters by (iii) evaluating high cell numbers per sample (200 000 cells) in a (iv) short time because samples are taken within generation times of bacteria (usually 60 min). The data should be available in-time to allow for on-line monitoring approaches.

## Previous work

To identify cell clusters, several approaches have been developed in the past three decades. These approaches can be classified into (i) manual, (ii) semiautomatic and (iii) fully automatic approaches.

i. Manual approaches are common and are represented by cytometric visualization and evaluation software like the commercially available `FCS Express` [18], the device-specific `FloMax` [19], and `Summit` [20], or the freeware `FlowPy` [21]. All of these tools provide a 2D graphical representation of cytometric data. The measured parameters (e.g. forward-scatter (FSC) or fluorescence intensity) used as axes of a 2D dot plot can be selected by the user. Each axis is divided into channels representing the signal intensity of an event after amplification. To mark cell clusters, the user can draw rectangle, ellipsoid, quadratic or polygonal regions inside the dot plot. Each of these regions identifies a cell cluster. The counts (number of cells) for each cell cluster can be extracted for further analysis. These approaches are time-consuming and user-dependent as the number of marked cell clusters as well as the position and the size of the marked regions is based on the experience of the user [22, 23, 24].
ii. Semiautomatic approaches are represented by cytometric visualization and evaluation software such as `FlowJo` [25]. Besides the manual marking of cell clusters, `FlowJo` provides a semiautomatic auto-clustering tool to identify cell clusters based on equal probability distributions which is restricting the shape of the clusters. The user can adjust the size and the shape of each identified cell cluster by moving the mouse over the dot plot and changing the vertices of the polygon gate. The number of clusters that can be identified in this way is not restricted and the counts for each cell cluster can also be extracted for further analysis. As the cell clusters are identified in a semiautomatic way, this approach is less time-consuming but the results still need manual effort by the user and are dependent on the user’s experience.
iii. Automated approaches comprise software tools that were developed to provide user-independent and reproducible clustering results of flow cytometry data. Recently, new approaches were developed to achieve clustering results automatically that fit the expectations of the user.

`flowFP` [26] is using the Probability Binning (PB) algorithm [27]. The binning procedure divides the two-dimensional dot plot into rectangular regions (bins) that contain nearly equal numbers of data points. This step of dividing the dot plot areas is performed multiple times based on the number of recursions adjusted by the user.

`SamSPECTRAL` [28] uses a modified spectral clustering algorithm which is based on data subsampling (faithful sampling), graph-theoretical principles and the k-Means algorithm [29]. `SamSPECTRAL` has the capability of identifying arbitrary shape clusters since it is a non-parametric approach that makes no assumptions on the shape and distribution of clusters. The main parameters of this approach are the scaling parameter *sigma* defining the “resolution” in the spectral clustering stage and the *separation factor* being a threshold that controls to what extend clusters should be combined or kept separated. In principle, with a larger *sigma* smaller clusters will be identified and with a larger *separation factor* more clusters will be identified. Both parameters have to be adjusted properly by the user. A strategy to adjust both parameters by reference to one’s own data is provided in the user manual of the package. The general way is to run `SamSPECTRAL` multiple times using the same data and to change both parameters until they fit the requirements described in the user manual.

The concept of `flowDensity` [30] is sequential bivariate clustering. `flowDensity` estimates the region around cell populations using characteristics of a marker density distribution (e.g. the number, height, and width of peaks and the slope of the distribution curve). Predefined cell subsets are identified based on the density distribution of the parent cell population by analyzing the peaks of the density curve. `flowDensity` aims to gate predefined cell populations of interest where the clustering strategy is known.

`flowMeans` [31] is based on the k-Means algorithm. The number of modes is counted in every single dimension followed by multidimensional clustering. Adjacent clusters are merged using Euclidean or Mahalanobis distance and the number of clusters is determined by a segmented regression algorithm to detect the change point in the distance between the merged clusters. As this approach is based on the k-Means algorithm which is not considering cluster distributions, it is used to find equal-sized, non-spherical clusters.

`flowClust` [24] is using a model-based clustering approach based on the estimation of distribution parameters of clusters by using the Expectation-Maximization (EM) algorithm. The number of clusters to be found can be fixed or determined by using the Bayesian Information Criterion (BIC) in a manual way. The number of data points per cluster is calculated and outliers can be identified by specifying quantiles (e.g. 90%) of the clusters. This approach is providing good results for Gaussian distributed cell clusters. An extension, `flowMerge` [32] provides automated selection of the best number of clusters, as well as merging overlapping cluster components.

`FLAME` [33] is an online software placed on the public server of the Broad Institute (Cambridge, Massachusetts, USA) and is a model-based clustering approach using the EM algorithm to estimate the distribution parameters of clusters. To determine the appropriate number of samples, the Scale-free Weighted Ratio (SWR) was invented. This measure is based on the average Mahalanobis distances, normalized for the distinct variances (which determine shape, dispersion, orientation, etc.) of different clusters, that are computed for pairs of points within and across clusters. `FLAME` also provides the construction of a global template of clusters which can be used to identify clusters across samples and to follow dynamics.

From this review, we can draw the conclusion that each of the stated tools has advantages towards manual clustering if the complexity of the data is not that high (e.g. a low number of clusters or a low number of data points) and the information content of the clustering results is restricted to general statements like the membership of a measured cell to one of the identified clusters. Each of these tools has different limitations. Based on the data we work with, and which forms the basis of the later evaluation, we point to the following limitations shared to some degree by the above-mentioned tools. Some of the tools are not practicable for microbial flow cytometry data that are different from medical data. The most important point here is the abundance of clusters we are faced with, while medical data tend to have few, mainly two, clusters in samples. Furthermore, our data have only two channels but a large number of distributed subcommunities within this range. We detect changes in the structure of bacterial communities by counting numbers of subcommunities per 2D-plot by recognizing the position of the subcommunities in the same 2D-plot and by counting cell numbers per subcommunity (technically, per gate). This type of analysis can be performed every few minutes without noticable effort time or financial effort. A wealth of information can be drawn from the clusters if community dynamics (i.e. dense sampling) are pursued.

Medical applications e.g. in oncology or hematology are the broadest field for the use of flow cytometry. Thus most of the automated approaches were designed to fit the requirements of these data sets. The cells measured in medical applications are relatively big (e.g. size of blood cells is around 10-20 *μ*m) and are usually labeled with differently fluorescent antibodies that specify the cell type. As a result, the cytometric data of one sample provide multiple fluorescence parameters besides the intrinsic cell parameters such as forward-scatter (FSC) or side-scatter (SSC). Several 2D plots are required to describe all cell types in a typical sample. Therefore, the number of gates per one 2D plot is frequently low and does not surpass 3 to 5 subpopulations which can be seen in data sets such as GvHD (graft-versus-host disease [34, 22], http://flowrepository.org/id/FR-FCM-ZZY2) or HSCT (hematopoietic stem cell transplant [22], http://flowrepository.org/id/FR-FCM-ZZY6).

In microbial applications, in particular in applications of microbial community analyses, the cells are much smaller (0.7-2 *μ*m) and are usually treated with only one fluorescent dye to mark all cells in a community and to separate the cells from noise and debris. Commonly, DAPI (4’,6-di-amidino-2-phenyl-indole) or SYBR Green are used that stain the DNA or the nucleic acid of all cells, respectively. In contrast to the highly resolving DAPI the resolution of microbial communities by SYBR Green is much lower and results mainly in only two subcommunities such as low nucleic acid (LNA) and high nucleic acid (HNA) bacteria. Recently, an attempt was made to resolve these two subcommunities even further by applying a deconvolution model [35].

Instead, the data generated from microbial community measurements using DAPI appear as highly complex system which encompass high numbers of taxonomic entities and fast variations in physiological states of the measured cells [4, 9, 10]. In addition, DAPI is prone to find rare cell types in a complex community as its fluorescence resolution is of high quality. As a result, the number of clusters within the cytometric dot plot can be very large and their separation becomes a difficult problem.

Based on our desiderata as mentioned above, this leads to the following requirements on the clustering algorithm: it must (i) be fast enough, (ii) determine the number of gates automatically, (iii) separate cell clusters from background clusters containing irrelevant information, and (iv) calculate the real number of data points for each cell cluster. The previously presented automated approaches hit their limits trying to by fulfill these requirements and do not produce adequate results. As a consequence, the identification of cell clusters is still performed in an experience-based, manual way in microbial flow cytometry. This severely limits the amount of data that can be processed. To improve this situation, we developed `flowEMMi`, a tool that is able to identify real cell cluster distributions in microbial FCM data quasi on-line in an automated way and to export necessary abundance information of every cell cluster for further analyses.

## Methods and implementation

### Conceptual Outline

Each single cell of a microbial community is visualized as a data point in a two dimensional cytometric dot plot. The cell is described by physiological properties such as cell size measured by forward-scatter (FSC) and number of chromosomes per cell measured by fluorescence intensity using DAPI (4’,6-di-amidino-2-phenylindole). Both physiological properties were used in recent studies of complex microbial community systems with success [36, 4, 9]. Additional parameters can also be used for evaluation such as cell density (side-scatter (SSC)) or pulse width [37, 38].

Cell clusters are typically drawn as ellipsoid regions within the dot plot by using cytometric visualization and evaluation software such as `Summit` or `FlowJo`. Ellipsoids as geometric boundaries make sense for at least three reasons. (i) They are easy to calculate. (ii) They conform to the way practitioners typically define boundaries of clusters in cytometry e.g. to define cell subsets for cell sorting. (iii) More importantly, an ellipsoidal shape conforms well enough to identified clusters in real data because cells typically distribute as bivariate Gaussian curves [37].

Let 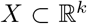 be the set of data points *x ∈ X* obtained from an experiment. The data considered here typically has *k* = 2, since clustering is performed on projections onto two parameters. Ellipsoid regions of arbitrary orientation are described via the equation (*x − υ*)^*T*^ **A**(*x − υ*) = 1 where *υ* is the vector-valued position of the center, **A** is a positive definite matrix, and *x* denotes solution vectors to the boundary. The corresponding statistical density function is the multivariate normal *P*(*x*) ∝ exp (−(*x − μ*)^*T*^Σ^−1^(*x − μ*)). Here 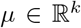 is the mean, 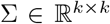 the covariance matrix, and 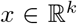 are points whose density is given by *P*(*x*). Having more than one Gaussian distribution leads to a mixture model ∑_*i*_ *π_i_P_i_*(*x*), with *π_i_* (*π_i_* ≥ 0, ∑ *π_i_* = 1) describing the weight/probability of each Gaussian. From a statistical point-of-view, multivariate normal distributions provide the framework with which to infer the most likely position of the ellipsoid regions [39, 40]. The parameter space of the model is written more succinctly as *θ* = (*π*, {*μ*_1_, …, *μ_n_*}, {Σ_1_, …, Σ_*n*_}) for a mixture model with *n* elements, hence log 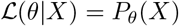.

As the real distribution parameters are unknown, all parameters, i.e., mean and covariance for each individual Gaussian and the weight vector *π* have to be estimated. Since no closed form solution exists, an iterative procedure has to be employed. It appears natural to use the expectation-maximization (EM, [41]) algorithm which is employed to find maximum likelihood estimates of unknown parameters of statistical models. The estimated parameters might not be the best solution as the EM algorithm is only guaranteed to converge to a *local* optimum.

In the E (expectation) step (Eqn. 1), the (log-)likelihood is calculated based on the estimated parameters of each cell cluster of the current iteration. In the M (maximization) step (Eqn. 2), new parameters of each cell cluster are computed to maximize the (log-) likelihood from the E step.

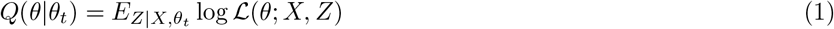

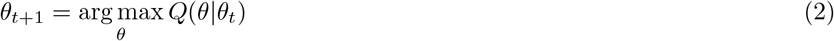

Both steps are performed iteratively until a termination condition is fulfilled by using the following criterion:

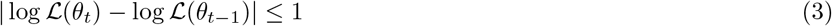

As the likelihood function is steadily growing, the estimated parameters of each cell cluster are converging toward a local optimum. Upon termination of the algorithm, every two-dimensional data point has a probability to belong to one of the determined cell clusters that are defined by the estimated parameters of the underlying distribution. Due to the steady growth of the log-likelihood function, the EM algorithm only finds one local optimum and has therefore to be initialized several times with different start values. Nevertheless, even after a large number of initializations it is possible that the global optimum for the numbers of initializations will not be found and that the calculated estimates of the parameters are not the best possible solution [42].

Usually, the EM algorithm needs to be initialized with start (prior) values for each distribution parameter in the first (E) step. The number of parameters *k* to be initialized is dependent on the number of clusters *c* and is equal to 6*c* − 1. For *c* = 20, the user would have to pick 119 start values that also need to fulfill some requirements (e.g. 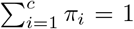). This is a time-consuming procedure and is prone to cause errors. For an easier initialization we changed the order of the steps by choosing the M step first. Thus, the probabilities of each data point belonging to one of each cluster are randomly sampled from a Dirichlet distribution, which can be used as a prior distribution for the probabilities [43] with hyper parameter α = 1 ensuring that the probabilities of one data point sum up to 1. Based on the probability matrix the distribution parameters are calculated first and in the next step the (log-) likelihood is calculated based on the estimates of the parameters of the first iteration. If good prior distribution parameters are available (after the subsampling procedure, see section Data reduction – Subsampling), these are used instead for the initialization.

### Implementation

To be able to pass objects from `R` to `C++` and back and to achieve an efficient implementation of the EM algorithm we used `Rcpp` [44] and the `Eigen C++` template library (version 3.3.3) which is provided by the `RcppEigen` package (version 0.3.3.3.1, [45]). As the EM algorithm is based on linear algebra operations, including matrix-vector and matrix-matrix operations, `RcppEigen` enables convenient access to a high-performance framework to implement these operations efficiently. This package needs to be installed to the `R` library and is essential for the use of `flowEMMi`.

Other packages that need to be installed to the `R` library for reading and working with the standardized `.fcs` files, visualizing the cytometric dot plots, calculating the statistical significance of the results and for the random initialization of the EM algorithm are `flowCore` [46], `flowViz` [47], `ggplot2` [48], `randomcoloR` [49], `mixtools` [50] and `gtools`[51].

### Removal of technical noise and beads

Technical noises (such as instrumental noise and cell debris) are unavoidable during a cytometric measurement. These are represented by extremely low fluorescence value or scatter signals in each dot plot. Before automatic determination of gates, such technical noises should be removed, in addition to the scatter and fluorescence signals of beads, which are implemented in each measurement for the alignment of samples. In this study, technical noises and beads per sample were removed with three steps (Figure 1) by the software `FlowJo` [25].

**Figure 1.**
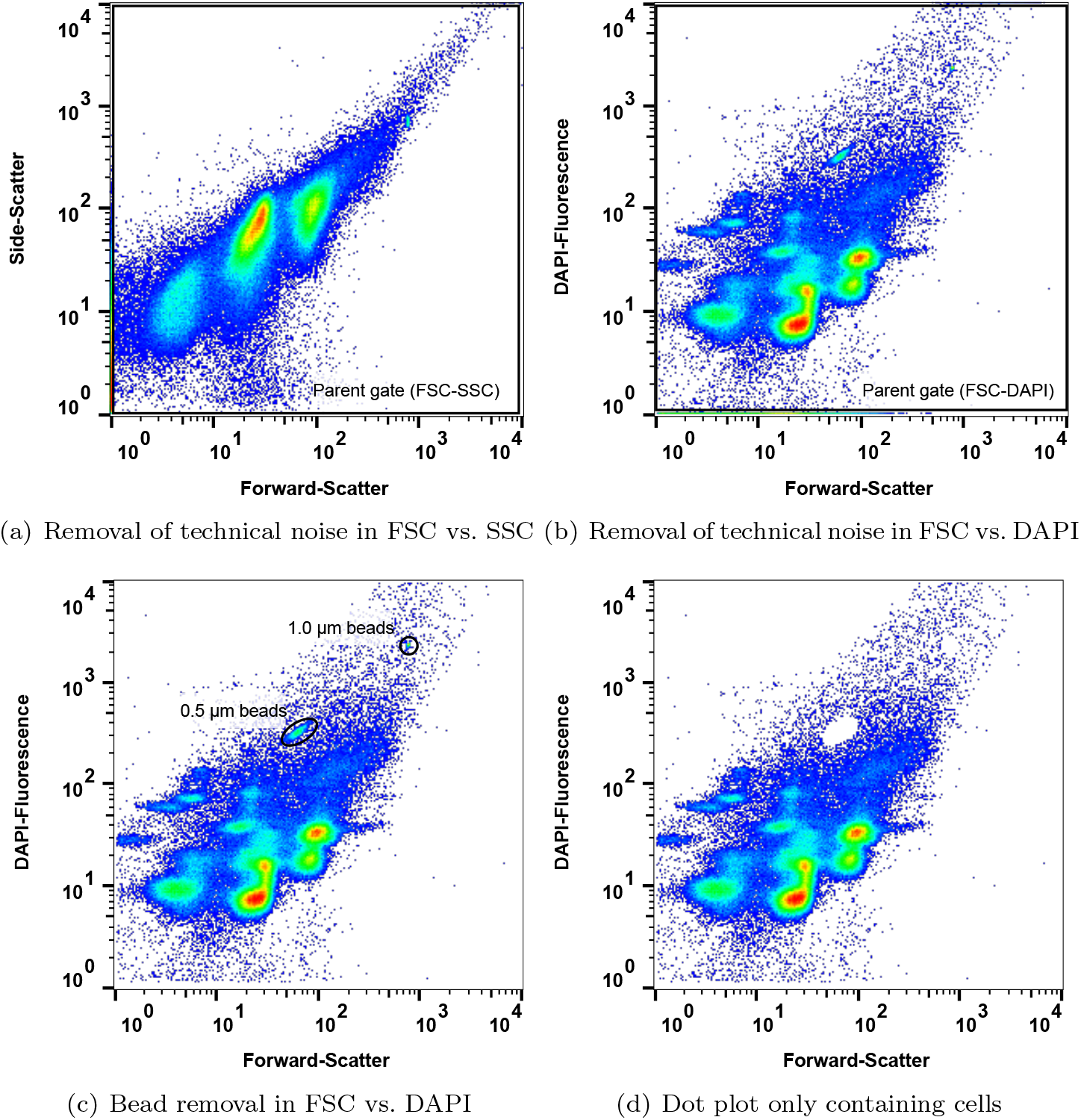
Steps for removing noise and beads from the cytometric dot plot. **(a)** Technical noise removed by setting a parent gate in forward-scatter vs. side-scatter. **(b)** Technical noise removed by setting a parent gate in forward-scatter vs. DAPI fluorescence. **(c)** Beads removed in forward-scatter vs. DAPI fluorescence. **(d)** Cytometric dot plot only containing cells used as input for `flowEMMi`.

First, all events were visualized in the 2D-dot plot of forward-scatter (FSC) vs. side-scatter (Figure 1, a), and a parent gate (FSC-SSC) was set to remove technical noises from FSC and SSC channels. Second, similarly, technical noise from the channel of the DAPI fluorescence was removed by setting a parent gate (FSC-DAPI) in the 2D-dot plot of FSC vs. DAPI fluorescence (Figure 1, b). Third, bead events were removed via specific gates (Figure 1, c) with the goal of retaining only events that represent cells (Figure 1, d). Once created, the `FlowJo` workspace containing all these steps can be saved and automatically applied to all samples of the experiment. The final data, only containing cell events, are used as input for `flowEMMi`.

### Finding the best number of clusters

In microbial flow cytometry a large number of clusters within one sample is very common. Furthermore, the actual number of clusters is unknown independent of the complexity of the data. To overcome the obstacle of a manual selection, `flowEMMi` was designed to determine this number automatically. Since the number of clusters is unknown, a (usually larger) range (e.g. *c* ∈ {2, …, 20}) has to be defined by the user to find all clusters at the first run of `flowEMMi`. A larger range is recommended because `flowEMMi` should generally have no parameters that need tuning and return the most appropriate number of clusters regardless of whether it is low or high. This prevents time-consuming initializations of the EM algorithm and an overestimation with excessive numbers of clusters.

To determine the most appropriate number of clusters we used the Bayesian Information Criterion (BIC, [52, 53]). Besides other model selection criteria like the Integrated Complete-data Likelihood (ICL) or Slope Heuristics, the BIC is known to provide the true number of clusters for Gaussian mixture models in most cases [39]. Such selection criteria have been used successfully before [32]. Eqn. 4 describes of calculation of the BIC for *c* clusters, with 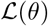 being the achieved likelihood for a model *θ* with *c* clusters, *k* parameters and *i* data points.

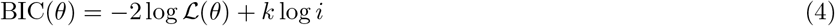

The BIC curve can be plotted and shows the BIC value for each number of clusters *c*. In most cases, the curve has a positive exponential trend and for a particular *c* the trend of the curve is getting nearly linear. Thus, the value of *c* at this point gives a good hint about the most appropriate number of clusters within the sample. Therefore, we defined a threshold for the difference of the BIC value between *c* and *c* + 1 for the whole range of *c*. If this difference is below 50 for the first time for the whole range of *c* then this particular value of *c* is considered as the most appropriate number of clusters *c*_BIC≤50_ for a given parameter set *θ*_*c*=1_, …, *θ*_*c=n*_ where we suppress the individual *θ*’s in the notation below:

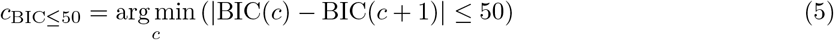

Nevertheless, it is possible to select a higher value for *c* than suggested by the BIC. A higher value would mean that more clusters are found but the increase of information/likelihood of the model is only marginal.

### Data reduction – Subsampling

The running time of the EM algorithm is dependent on the dimension of the data points, the initialization values of the model parameters (*π, μ*, Σ), the number of data points *i* and the number of clusters *c* to be found. The dimension is constant and as the initialization values are sampled randomly (see section Idea, last paragraph), the effect on the running time of the EM cannot be determined or adjusted from the very start. Good initialization values result in a low number of iterations and therefore decrease the running time whereby poor initialization values result in a high number of iterations and therefore increase the running time.

The number of data points *i* used for the clustering can be adjusted and has a notable and measurable influence on the running time. It was shown before that the results of the EM running on a subset of all data points is likely to provide distribution parameters that differ not that much from the distribution parameters resulting from the EM running on all data points [54]. As the measurement of the cells does not follow a certain order (e.g. small cells first, big cells last), the data used as input for the EM are unordered, too.

Thus, a subset of cells can easily be selected by choosing e.g. every 20^*th*^ data point of the full data set. With the selection of a subset it is possible to reduce the running time of the EM in order to rapidly get a good approximation for the estimates of the model parameters (*π, μ*, Σ) of each cluster *c*. For the evaluation of `flowEMMi` we used samples containing 200 000 cells (without noise and beads) which ensures a high statistical significance of the appearance of cells in respective segregated subsets. Measuring fewer cells produces less precise statistical data, therefore, subsampling is recommended instead of working with fewer measured cell numbers per sample. By creating a subset with, say, every 20^*th*^ data point big clusters will still be visible and detected by `flowEMMi`. Only those clusters with a very low abundance may get lost. By combining the subsampling procedure with the BIC (see section Finding the best number of clusters), the best number of clusters *c* can also be determined automatically in a very short time.

Consequently, the reduction of the number of data points and the use of the BIC reduces the running time of the EM and provides the most appropriate value for the number of clusters *c* as well as estimates for the model parameters of each cluster (*π, μ*, Σ). After this step, these outcomes can be used as already fitted initialization values for the EM running on the full data set thus preventing an elaborate and inaccurate initialization. In addition, instead of random initialization values for samples with similar structures the same fitted initialization values can be used as input which further increases comparison between samples and decreases the running time substantially.

### Data separation

Another important step is to eliminate irrelevant data points occurring from technical noise, beads or cell debris. These data points are not needed for the analysis of the cell clusters and therefore have to be separated from the real data representing the cells (see section Removal of technical noise and beads). In addition, not all cells cluster as condensed ellipsoid regions and are instead more evenly distributed across the dot plot. As every cluster algorithm generally is designed to assign every data point to one cluster, a mixed model was developed to create a background model for the evenly distributed data points and a foreground model for the relevant cell data points.

Cell clusters form condensed ellipsoid regions within the dot plot but the data points of a background cluster spread over a large area. Thus, the variance (of the main diagonal of) Σ of a cell cluster distribution is much smaller than the variance of a background cluster distribution. A threshold can be defined to separate the clusters with very high variance from the clusters with smaller variance. A maximum standard deviation *σ* (square root of variance) value is predefined to separate background clusters from foreground clusters but can be changed by the user if required e.g. if only very small clusters (rare subcommunities) or bigger clusters (dominant subcommunities) should be found. We default to a setting where a cluster is set as a background cluster if min_*d*_(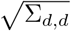) ≥ 2500.

Nevertheless, it was shown that the cell numbers of background distributions, denoted as off-gate cells, are of importance as they can be an indicator for occurring disturbances in microbial systems [4]. For this reason, the off-gate cell number of all background clusters is saved in a readable text file besides all cell numbers of the detected cell clusters and can be used for further analyses.

### Calculation of cell numbers/Confidence intervals

After the clustering procedure, the reduction of the data set, and the separation of background clusters, some cell clusters may not have a clear ellipsoid shape and can also contain outliers. For Gaussian distributions, the calculation of confidence intervals [55] is a statistically legitimate way to select data points having a certain significance for being part of an identified cluster. As a confidence interval is the complement of the level of significance (usually called p-value), a 95% confidence interval reflects a significance level of 0.05 which is most commonly used in statistics [56].

For multivariate Gaussian distributions the confidence interval of each cell cluster *c* is determined by its mean vector *μ* and its covariance matrix Σ. Based on these distribution parameters, the data points lying inside the confidence interval *q* = 1− *p* can be calculated using the following equation.

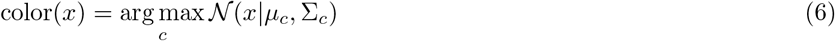

A point *x* is considered to be part of the confidence interval for the color *c*, if the following two Dirac-*δ*-functions determine that both, the optimal color for *x* is *c*, and the scaled density is higher than *q*.

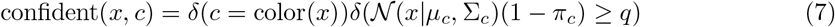

As a consequence, the shape of the data points enclosed by the confidence interval is elliptic. If required, the user can change the confidence level (e.g. to 90% or 99%). After the calculation of the confidence intervals, the numbers of data points of each cell cluster are saved in a `.txt` file. These numbers can be used to compare the result of the automated clustering with the results of the manual clustering and for further analyses.

## Results and conclusion

A real FCM sample set containing 10 samples (https://flowrepository.org/id/RvFrxWfLS98ghWy8f9bAx7E0JWpcgNiIfpBmJjtbyScv1SgtgF8ACPnlXGEkTHYb) was used to investigate if the methods described above provide adequate results and make our tool `flowEMMi` suitable for automatic clustering in cytometric microbial community data. In this section, the sample InTH_160712_025.fcs (Figure 1) was used representatively. The clustering results of the other 9 samples can be found in the Supplementary information (supplementary Figures 5–14).

First, we tested whether the optimal number of clusters *c* can be determined using the BIC. To achieve good approximations for the number of clusters *c* as well as for the estimates of the model parameters of each cluster (*π, μ*, Σ) in this sample, the EM algorithm was initialized with a subsample of all data points (every 40^*th*^ data point) and randomly sampled cluster probabilities for each data point. Before subsampling, technical noise and beads were removed. After removing noise and beads 200 000 data points remained in the `.fcs` file. As only every 40^th^ data point was used in this step this means that only 5000 data points were used as input of `flowEMMi`. As the real number of clusters *c* was unknown the minimum number of clusters to be found was set to 2 and the maximum number of clusters to be found was set to 20. Figure 2 shows the results of `flowEMMi` after subsampling and calculation of the BIC.

To show the impact on the choice of the number of clusters we provide, in addition to the estimated number of *c* = 13 clusters (Figure 2c), two extra clustering results. One result gave a very low number of *c* = 5 clusters (Figure 2d), and the other of *c* = 20 which are too many clusters (Figure 2e). *c* = 13 seems to be an appropriate value as the trend of the BIC curve is nearly linear after this point. The difference of the BIC value between *c* = 13 and *c* = 14 is below 50 for the first time for the whole range of *c*. This result can be derived by looking at the plot of the BIC curve (Figure 2a) as well as output information of `flowEMMi`.

As the number of data points in the subsampling step is only 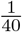 of the original number it is possible that some clusters are missed after subsampling. Due to that and the possibility that one or more clusters may be identified as background clusters in the full data run, *c* = 13 can be seen as a very conservative value which allows to identify the main clusters. If the user wants to detect rare clusters the number of clusters to be found should be set higher than suggested by the BIC for the full data run (e.g. *c* ∈ {13, …, 16}). We can say that the first run of `flowEMMi` using a subsampled data set and the BIC provided the appropriate value for the number of clusters *c* as shown by the BIC curve (Figure 2a) and good approximations for the parameter estimates (*π, μ*, Σ) of each cluster. These data are saved as output of `flowEMMi` and can be used as prior parameters for the full data run.

Then it was tested if the subsampling procedure decreases the running time by providing good estimates for the number of clusters *c* and good approximations for the parameter estimates of each cluster. `flowEMMi` was executed three times without and with usage of the subsampling procedure. Both ways were compared by measuring the total running times as well as the numbers of iterations for each of the three runs needed for *c* ∈ {13, …, 16}. Without subsampling `flowEMMi` was initialized with a range for the number of clusters to find *c* ∈ {2, …, 20} with only one initialization. With subsampling, the same range was defined and 10 random initializations were executed. Then, the outputs of the subsampling procedure were used as input for the full data run with a smaller range for *c* ∈ {13, …, 16} and only one initialization was executed to keep the comparability to the values achieved without subsampling. Table 1 shows the outcomes of this comparison.

**Table 1.**
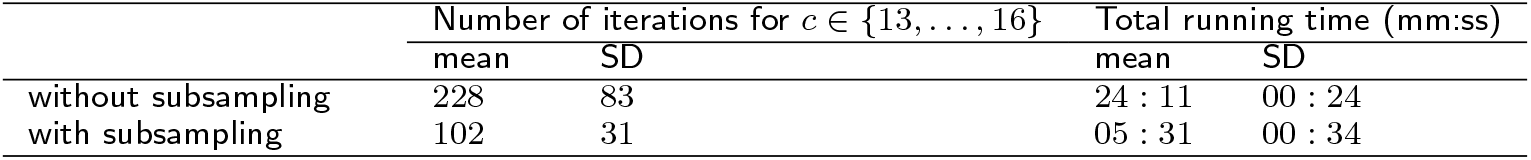
Comparison of running time without and with usage of the subsampling procedure. Mean values (mean) and standard deviations (SD) of the total running time and the number of iterations for *c* ∈ {13, …, 16} were calculated based on three executions of `flowEMMi`, respectively.

We can draw the conclusion that the subsampling procedure decreases the number of iterations needed for *c* ∈ {13, …, 16} up to approximately 60% with a much smaller standard deviation and the total running time up to approximately 75% with a nearly similar standard deviation. Therefore, we strongly recommend to use the subsampling procedure in order to achieve good results in a short time.

In the next step, the full data set is used as input with the extended range for *c* ∈ {13 … 16} to find rare clusters as calculated by the BIC and the prior parameter estimates of each cluster as calculated before by use of the subsampling procedure. Now, an additional threshold was defined to separate cell clusters from background. Figure 3 shows the final clustering results of `flowEMMi` running on the full data set.

Only clusters with a standard deviation below the threshold value are marked as cell clusters and are plotted in colors distinct from the gray background. In the next section, a benchmark procedure is performed to compare these final results i) to the results of manual clustering using `FlowJo` and ii) to the results obtained by the other tools.

### Benchmarking

To compare the results of `flowEMMi` with the manual clustering procedure, the sample In1TH_160712_025 (Figure 1, 2 and 3) was clustered independently by five expert users to identify the number of clusters, the range of the abundance values of all clusters and the percentage of background and foreground cell numbers based on 200 000 cells. For manual clustering the commercial program `FlowJo` was used and for comparison with the data generated by `flowEMMi` the data were biexponential transformed as by default. the following formula was used: 10^(mean/(65 536/4))^. Note: The value of 65 536 corresponds to the resolution of the cytometer device (here: InFlux, BD Bioscience, New Jersey, USA). For comparison, only those clusters found by `flowEMMi` that have the same or similar mean values in both parameters (FSC and DAPI-Fluorescence) as the clusters found by manual clustering are considered and counted. Table 2 shows the outcome of this comparison. The mean values of each cluster calculated by `FlowJo` and `flowEMMi` are given in the additional file `025.csv` which is part of the file `tables.tar.gz`.

**Table 2.** Comparison of clustering results from manual clustering performed by 5 experts using FlowJo and automated clustering using `flowEMMi`. Compared were i) the number of clusters that were found, ii) the range of the abundance values of all clusters, iii) the cell numbers of foreground/background cell and iv) the number of congruent clusters that were found by the user and `flowEMMi`, respectively. Congruent clusters are cell clusters having the same or similar mean values in both parameters (FSC and DAPI-Fluorescence).

|  | # clusters | Range of abundances (%) | Cell numbers (%) |  | # congruent clusters |
| --- | --- | --- | --- | --- | --- |
|  |  |  | Foreground | Background |  |
| flowEMMi | 12 | 1.56 - 20.72 | 71.6 | 28.4 | 12 |
| User 1 | 13 | 0.25 - 27.7 | 76.5 | 23.5 | 10 |
| User 2 | 15 | 0.21 - 30.0 | 82.1 | 17.9 | 11 |
| User 3 | 13 | 0.24 - 28.2 | 79.1 | 20.9 | 11 |
| User 4 | 16 | 0.26 - 31.2 | 90.7 | 9.3 | 11 |
| User 5 | 15 | 0.22 - 32.1 | 91.6 | 8.4 | 11 |

It can be seen that the number of cell clusters found by the expert users is in the similar range of the 12 clusters found by `flowEMMi` and varying from 13 to 16. The range of abundances and the proportions of foreground and background cell numbers is slightly in favor of the expert users which covered more cells within the clusters. Further 9 samples were tested in this regard. The outcomes can be seen in the Supplementary information. For `flowEMMi` and all previously introduced automated clustering tools (`flowFP, SamSPECTRAL, flowDensity, flowMeans, flowClust` and `FLAME`) we also compared the running time, the abilities to determine the number of cell clusters automatically, to separate cell clusters from background clusters and to calculate the cell numbers for each cell cluster. Table 3 shows the results of this comparison.

**Table 3.** Comparison of automated clustering approaches. Automated approaches were compared regarding the running time and the abilities to identify rare cell types, to separate cell clusters from background clusters and to calculate the real cell numbers for each cell cluster. ^*a*^ Running time calculated on a Intel(R) Core(TM) i5-3210M CPU @ 2.5 GHz with 4096MB RAM and Windows 7 Enterprise 64-Bit Edition. `FLAME`: “−*“ denotes that no results were received as our submitted “jobs” were always in the queue for several days and later cancelled by the server. `flowEMMi` is the implementation discussed in this work. *space-part*. denotes k-means type algorithms that do not produce tight clusters.

| Tool | Running time (h:mm:ss)<sup>a</sup> | Output features |  |  |  |
| --- | --- | --- | --- | --- | --- |
|  |  | Determine number of clusters | Shape of clusters | Separate background | Calculate cell numbers |
| flowEMMi | 0 : 05 : 31 | yes | ellipsoid | yes | yes |
| flowFP | 0 : 00 : 03 | no | rectangular | no | not applicable |
| SamSPECTRAL | 0 : 06 : 25 | space-part. | arbitrary | no | not applicable |
| flowDensity | 0 : 00 : 02 | no | rectangular | no | not applicable |
| flowMeans | 0 : 00 : 17 | space-part. | non-spherical | no | not applicable |
| flowClust | 1 : 15 : 30 | yes | ellipsoid | no | yes |
| flowMerge | (Table 4) | yes | ellipsoid | yes | yes |
| FLAME | —* | —* | —* | —* | —* |

In addition to this table, the clustering results of all tools are displayed in Figure 4. Results for `flowMerge` have been separated out into Table 4 and supplementary Figure 15, as `flowMerge` has a similar feature set. From a user standpoint, the better F_1_ measure and vastly improved running times of `flowEMMi` are most important.

**Table 4.**
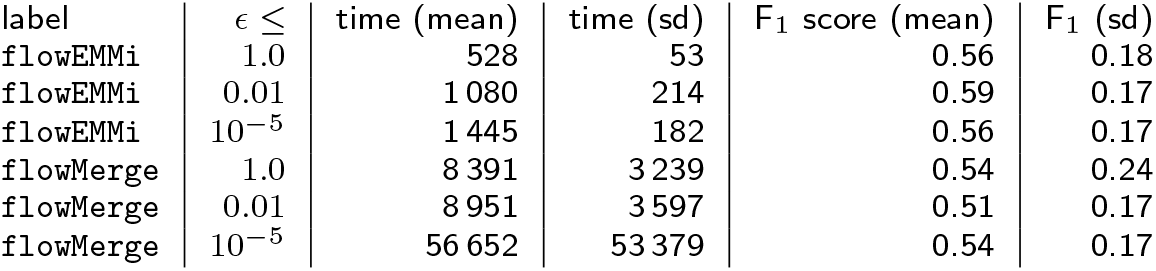
Running times and F_1_ score aggregated over experiments with different *ϵ* stopping criteria. Times and F_1_ scores (and their standard deviation (sd)) are aggregated over four experiments and 5 expert user gatings, each. Note that the default `flowMerge` stopping criterion of 10^−5^ yields running times in excess of 1 day. `flowEMMi` consistently yields better F_1_ measures with an average improvement of 4% to 16% over `flowMerge`, with much better running times, easily yielding speed improvements of ×8 − ×15 or better. For both algorithms, having a more stringent EM stopping criterion tends to increase the F_1_ score, however especially for `flowMerge` at prohibitive running time costs.

| label | $\epsilon \leq$ | time (mean) | time (sd) | $F_1$ score (mean) | $F_1$ (sd) |
| --- | --- | --- | --- | --- | --- |
| flowEMMi | 1.0 | 528 | 53 | 0.56 | 0.18 |
| flowEMMi | 0.01 | 1 080 | 214 | 0.59 | 0.17 |
| flowEMMi | $10^{-5}$ | 1 445 | 182 | 0.56 | 0.17 |
| flowMerge | 1.0 | 8 391 | 3 239 | 0.54 | 0.24 |
| flowMerge | 0.01 | 8 951 | 3 597 | 0.51 | 0.17 |
| flowMerge | $10^{-5}$ | 56 652 | 53 379 | 0.54 | 0.17 |

Table 3 and Figure 4 show that none of the other tools used for this comparison can separate cell clusters from background clusters. This is important as not only the cell numbers of real cell clusters can be used for further evaluation tools but also background cells as additional information which is useful for some applications. Both, the number of predicted cell clusters, and the relative number of cells, predicted to be part of clusters are important. As such, any tool that severely underestimates the number of clusters will almost certainly mispredict the number of cells that form clusters which makes downstream analysis complicated and can lead to misinterpretation of results. A reasonable estimate of the correct number of clusters is only provided by `flowEMMi, flowClust`, and `flowMerge`. For `SamSPECTRAL` and `flowMeans` the calculated number is in a good range but too small. The correct shape of clusters and therefore the required distribution parameters are only provided by `flowEMMi, flowClust` and `SamSPECTRAL`. The shape of clusters produced by all other tools does not reflect real cell cluster distributions. In addition, the running time of `flowEMMi` is with 15 times lower than the running time of `flowClust` and achieves better results. The `flowMerge` software extends `flowClust` and is the most close in terms of the set of features we require and therefore we performed an in-depth analysis of its F_1_ measure and running times compared to `flowEMMi`. In comparison, as shown in Table 4, `flowEMMi` delivers higher F_1_ values at vastly superior running times.

`flowEMMi` provides all information needed for the evaluation of microbial community FCM data. It fulfills all the requirements of the users and outperforms other tools that were tested with regard to running time and output features.

## Mock Communities

For additional testing of `flowEMMi` two artificial microbial cytometric mock communities were used consisting of either three or four different bacterial species. One of the artificial communities was harvested from plates and comprised three strains, namely *Stenotrophomonas rhizophila* DSM 14405, Kocuria rhizophila DSM 348, and *Paenibacillus polymyxa* DSM 36, while the other was harvested from liquid culture and comprised four strains, namely *Stenotrophomonas rhizophila* DSM 14405, *Escherichia coli* DSM 4230, *Kocuria rhizophila* DSM 348, and *Paenibacillus polymyxa* DSM 36.

The respective strains were separately cultivated in in Lysogeny Broth (LB, composition: yeast extract 5gl^−1^, NaCl 5gl^−1^, tryptone 10gl^−1^, pH 7.0, and agar 20 gl^−1^ in case of plates; Carl Roth GmbH, Karlsruhe, Germany). The cells were harvested and washed in PBS as described elsewhere [36], stabilized by adding para-formaldehyde solution (PFA, 2% in PBS), and, after a washing step, fixed in ethanol (70% in bi-distilled water) for storage at −20°C. DNA staining was performed using DAPI as described by [36]. For the plate microbial cytometric mock community the strains were mixed at proportions: *S. rhizophila*: 1%, *K. rhizophila*: 19%, *P. polymyxa*: 80%; and for the liquid microbial cytometric mock community the strains were mixed at proportions *S. rhizophila*: 2.5%; *K. rhizophila*: 20%, *P. polymyxa*: 70 %, and *E. coli* 7.5% as was determined by OD (d_λ 700nm_ = 0.5 cm). Finally, the two mock communities were measured by flow cytometry with regard to blue fluorescence (355 nm excitation) vs. forward scatter (488 nm excitation) using a BD Influx v7 Cell Sorter (Becton, Dickinson and Company, Franklin Lakes, NJ, USA). The raw data are available at https://flowrepository.org/id/RvFrfbZjwuR0FdR6H12EKWkDFj4CHNpRejkp2Hn2qFkR144zYbuwSB8AuzET6xXZ (file plate mock community: `mCMC80.1.19.fcs`; file liquid mock community: `70.2.5.20.7.5.fcs`).

The resulting flow cytometric patterns are shown in supplementary Figure 16 while the results of `flowEMMi` are shown in supplementary Figures 17 and 18. The data clearly show the powerful performance of `flowEMMi` which not only could separate the four, respective three strains of the two microbial cytometric mock communities but also even subpopulations of the used pure mock strains.

## Discussion

We compared the outcomes of `flowEMMi` to the outcomes of the manual clustering performed by 5 expert users (Table 2) using `FlowJo` based on one representative sample (Figure 1, 2 and 3). The clusters found manually by using `FlowJo` and automatically by `flowEMMi` were very similar concerning the percental abundances and the location of the cell clusters. `flowEMMi` slightly underestimated the abundances of cell clusters which might be caused by the fact that manually set clusters do not follow statistical conditions e.g. confidence intervals. Cell clusters only containing a small number of cells typically (at least for our data) do not conform to a Gaussian distribution, and instead have a mostly flat density.

Furthermore, cell clusters that are big and isolated very often vary in size and comprise only low numbers of cells which nevertheless seem to belong to the respective cluster but without statistical confidence. `flowEMMi` may not recognize such clusters since the cells might not be within the required confidence interval of the respective cluster and thus are not assigned with statistical significance. This gives an additional value to the quality of the clustering result. Nevertheless, the size of the cell clusters can be controlled by the user by decreasing or increasing the confidence interval.

We put an additional focus on the comparisons between `flowEMMi` and other automated approaches (Table 3). By using `flowFP`, one bin is always divided into two smaller bins of the same size. Therefore, the size and the location of each cluster is constrained to spatial subdivison and the number of clusters to be found is always a power of two where the exponent is the recursion depth. The clustering results can also not be used for cell sorting as the cells of interest are always surrounded by a rectangular region that contains more cells which are not of interest and is not reflecting the real distribution of the cell cluster.

By using `SamSPECTRAL`, even with adjustment of both parameters (*sigma* and *separation factor*) as described in the user manual, the number of clusters that were found was in general too small. Besides, the final results of `SamSPECTRAL` are always achieved after a subsampling procedure which is necessary to keep the running time of large data sets in an acceptable scale. The cell numbers per cluster are therefore always relative to the numbers of cells of the reduced input data. `flowDensity` is primarily designed to gate predefined cell populations of interest where the clustering strategy is known. As densities of cell clusters are often overlapping within one parameter (clusters with similar forward-scatter, i.e. cell size but different fluorescence intensity, i.e. number of chromosomes), these overlapping densities conflate into one big density distribution with one very wide peak what makes the separation almost impossible. Therefore, this approach is only suitable if the cluster densities are not overlapping to a high extend.

`flowMeans` is designed to find equal-sized, non-spherical clusters. Therefore, this approach is not suitable for Gaussian distributed clusters that form ellipsoid shapes and are very diverse in size. By using `flowClust`, background clusters that are evenly distributed across the dot plot are not separated from cell clusters. Besides, the running time of `flowClust` is relatively long and the number of cell clusters that are found is too low. We were not able to receive results from the online tool `FLAME` as our submitted “jobs” were always in the queue for several days and later cancelled by the server.

To increase the reliability of finding correct clusters concerning the location and abundances of cells, we used a model-based approach to determine the parameters of a mixture of multivariate Gaussian distributions. Our current implementation of the EM algorithm utilizes a variant of stochastic EM, which initializes the EM with different starting points for each run. Naturally, this will lead to slightly different clustering results for each run. Despite the fundamental assumption that cells form Gaussian distributed clusters it is also possible that different cell cluster distributions occur (e.g. flat distributions). In general, the EM algorithm is able to estimate the parameters of each existing distribution and also mixtures of different distributions. It is possible to fit parameters of different distributions to each cluster and to select which distribution is describing the cluster more precisely from a statistical point of view. We focused here on Gaussian distributions and achieved satisfactory results. Allowing different distributions could lead to better results as also cell cluster would be found that occur as e.g. clusters with essentially flat densities.

## Conclusion & Outlook

In this work, we devised a method for the automated clustering of flow cytometry data derived from microbial communities. There is a big demand for an automated clustering procedure for the evaluation of cytometric samples derived from biotechnology, natural environment as well as agricultural und human health disciplines e.g. the animal or human microbiomes [57].

Flow cytometric analysis of microbial communities were recently proven to provide much deeper insight into underlying mechanisms of community assembly in comparison to amplicon sequencing technologies [8]. Resolving the respective contributions of e.g. deterministic or neutral paradigms to community structure and functions is dependent on sample density which cannot be provided by any other method within community observation time. Thus, the automated clustering procedure derived from microbial communities contributes to an even faster evaluation procedure and would close a gap in currently available automated clustering procedures that were mainly developed for samples with eukaryotic background and diversification in many fluorescent channels thus providing only few subpopulations per 2D dot plot.

Our automated procedure is now able to find a high number of previously unknown distributions in one 2D dot plot which is a huge step forward for fast and nearly on-line disposal of data to allow interventions for process control or fast diagnostic decisions. Follow up tools for on-line data evaluation were recently published [7].

As cell clusters can not always be described as Gaussian distributions the next step will be to allow different types of distributions (e.g. distributions with flat densities) and to fit the most probable distribution to each cluster. This will allow `flowEMMi` to find more clusters being better described by the underlying distribution. The EM algorithm is a powerful approach to estimate the unknown parameters of distributions describing clusters of cells with equal or similar optical parameters that are measured by FCM. With this approach it is possible to overcome the user-dependent and time-consuming clustering procedure which is still performed in a manual way.

## Abbreviations

BIC: Bayesian information criterion
DAPI: 4’,6-di-amidino-2-phenyl-indole
EM: Expectation maximization
FCM: Flow cytometry
FSC: Forward-scatter
GvHD: Graft-versus-host disease
HNA: Iigh nucleic acid
HSCT: Iematopoietic stem cell transplant
LNA: Low nucleic acid
PB: Probability binning
SD: Standard deviation
SSC: Side-scatter
SWR: Scale-free weighted ration
SYBR Green: N’,N’-dimethyl-N-[4-[(E)-(3-methyl-1,3-benzothiazol-2-ylidene)methyl]-1-phenylquinolin-1-ium-2-yl]-N-propylpropane-1,3-diamine

## Declarations

### Author’s contributions

J.L. and C.H.z.S. developed the flowEMMi algorithm and wrote the implementation, Z.L. performed the experiments; J.L., S.M. and C.H.z.S. designed the research and J.L., S.M., C.H.z.S. and P.F.S. wrote the manuscript. All authors contributed critically to the drafts and gave final approval for publication.

### Availability of data and materials

The `flowEMMi` sources and additional files are available here: http://www.bioinf.uni-leipzig.de/Software/flowEMMi/.

## Acknowledgements

The authors thank Thomas Hübschmann, Florian Schattenberg, Susanne Günther and Johannes Lambrecht for performing the manual clustering with `FlowJo`.

## Ethics approval and consent to participate

not applicable

## Consent to publish

not applicable

## Supplementary information

**Figure 2.**
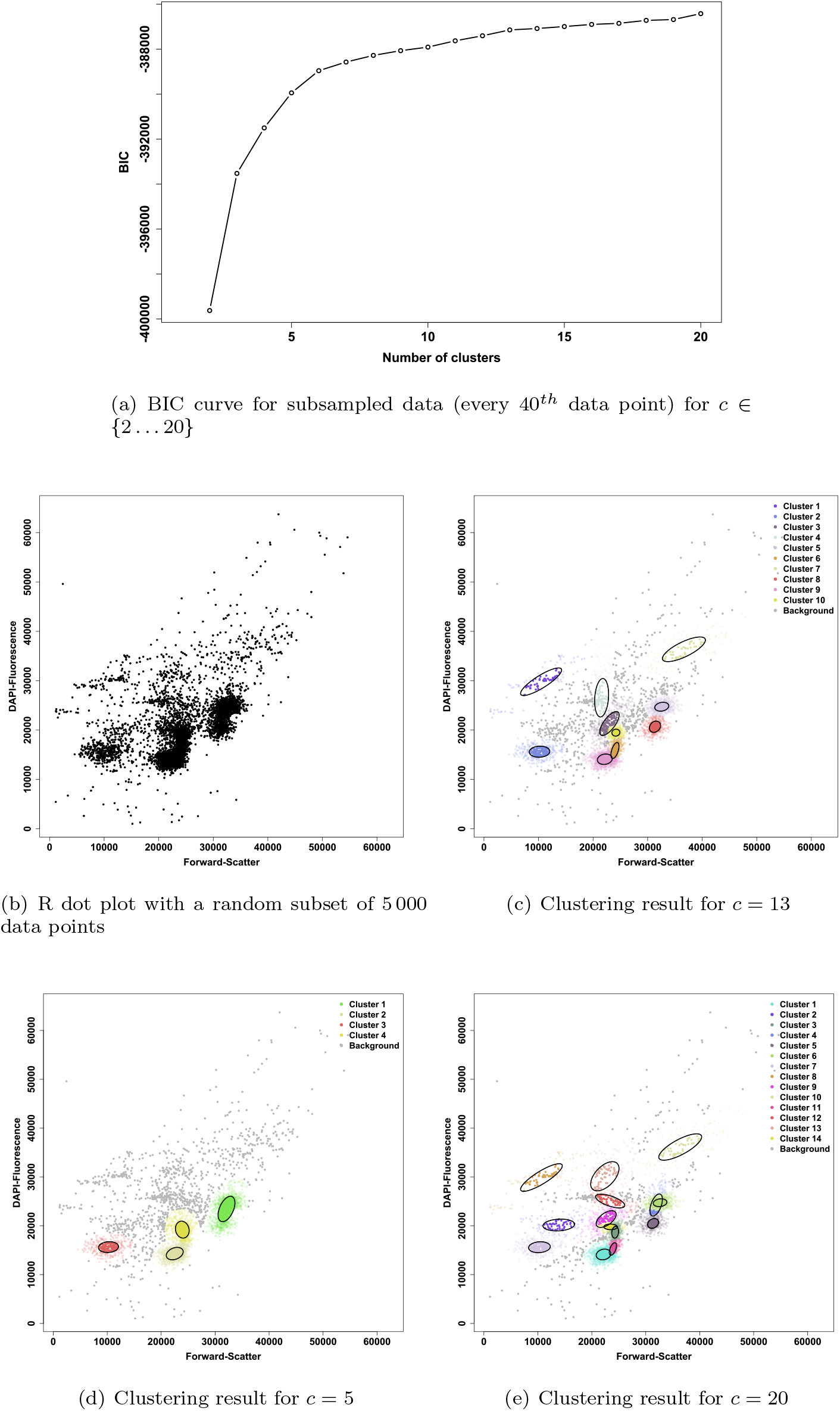
Results of `flowEMMi` after subsampling and calculation of the BIC for the sample shown in Figure 1 with separation of cell clusters and background clusters. Background clusters are not encircled and have a gray colour. **(a)** Curve of the BIC value shown for *c* ∈ {2 … 20}. **(b)** `R` dot plot with linear axes values from 0 to 65536 containing only every 40^th^ data point. **(c)** Clustering result of `flowEMMi` for *c* = 13 calculated as the most appropriate number of clusters with 10 cell clusters and 3 background clusters. **(d)** Clustering result of `flowEMMi` for *c* = 5 with 4 cell clusters and 1 background cluster. **(e)** Clustering result of `flowEMMi` for *c* = 20 with 14 cell clusters and 6 background clusters.

**Figure 3.**
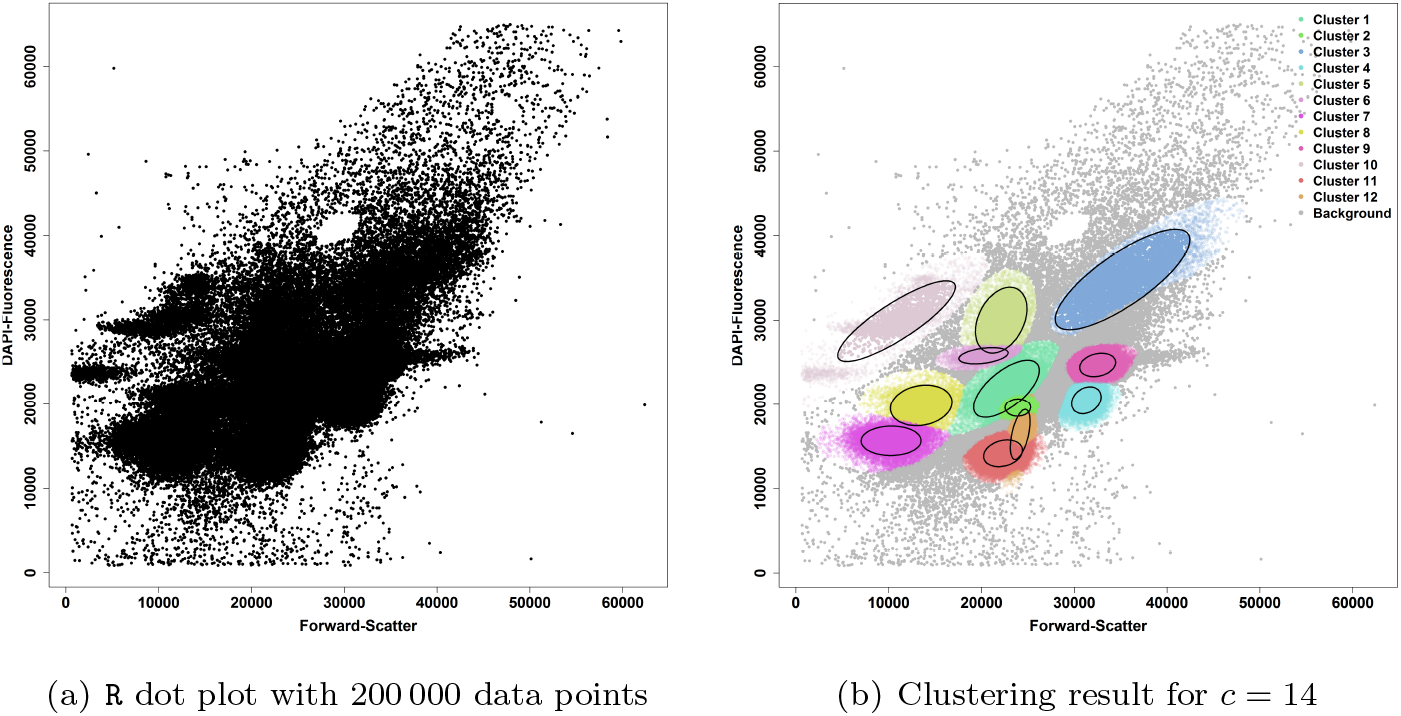
Final result of `flowEMMi` using prior distribution parameters achieved from the subsamling procedure and an extended range of *c* ∈ {13 … 16} achieved from the BIC to find rare cell clusters. **(a)** `R` dot plot with linear axes values from 0 to 65 536 containing all data points. **(b)** Clustering result of `flowEMMi` for *c* = 14 with 12 cell clusters and 2 background clusters. Background clusters are not encircled and have a gray colour.

**Figure 4.**
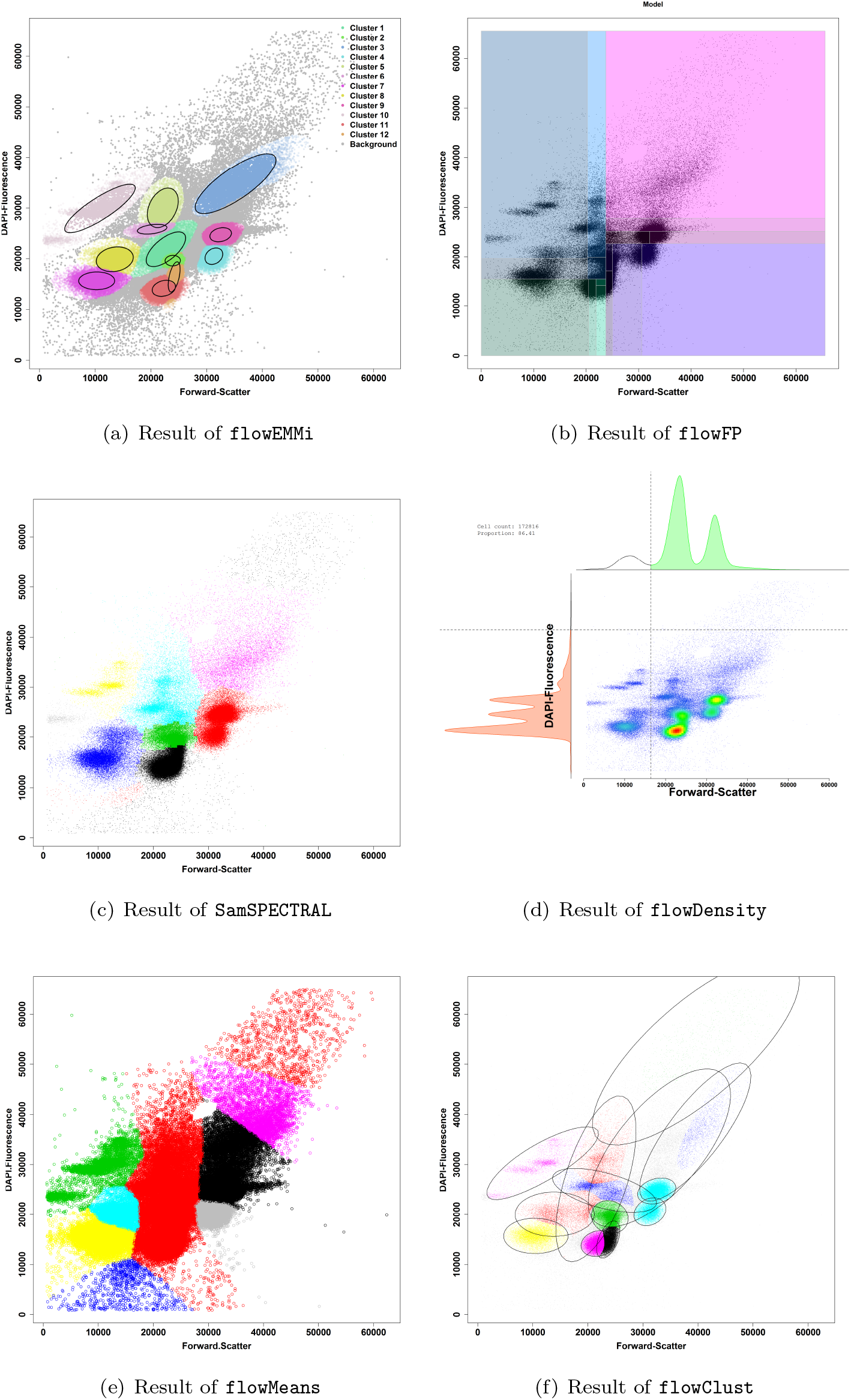
Results of clustering tools. **(a)** Result of `flowEMMi`. 12 cell clusters and 2 background cluster were identified. **(b)** Result of `flowFP` for 4 recursion = 16 clusters. **(c)** Result for `SamSPECTRAL` with adjusted parameters (*σ* = 1 000, separation=0.3) and automatically determined number of clusters. **(d)** Result of flowDensity with overlapping densities. **(e)** Result of flowMeans with Voronoi like cluster shapes (MaxN=20). **(f)** Result of flowClust with automatically determined best number of clusters for *c* ∈ {2 … 20} (cf. detailed analysis of `flowMerge` in Table 4 and discussion).

**Figure 5.**
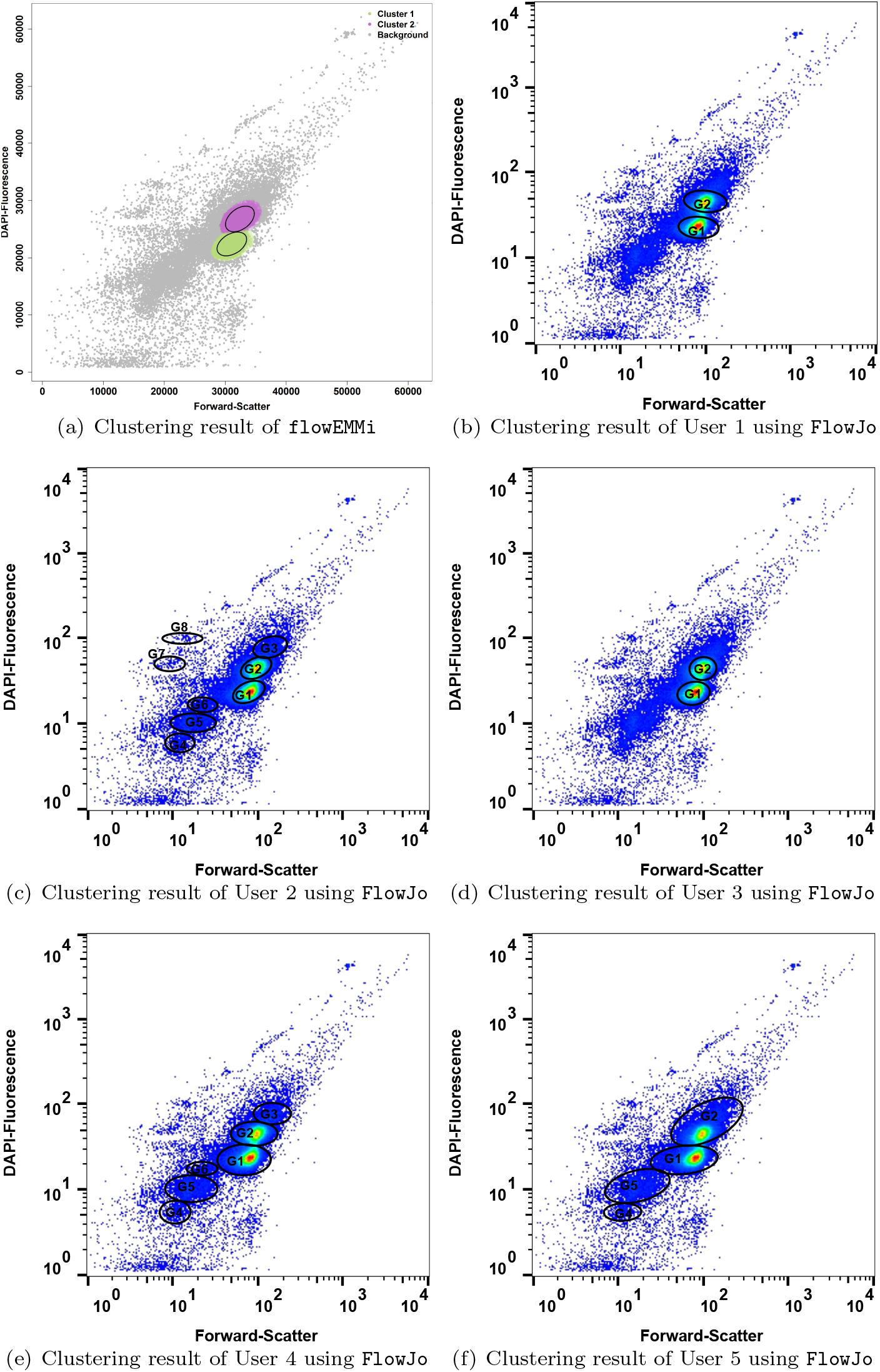
Clustering results for sample InTH_160719 039 using `flowEMMi` with 2 congruent cell clusters and 94.1% foreground cells **(a)** and manual clustering performed by 5 expert users using `FlowJo` **(b-f)**. User 1 selected 2 cell clusters with 89.9% foreground cells **(b)**. User 2 selected 8 cell clusters with 93.4% foreground cells **(c)**. User 3 selected 2 cell clusters with 91.1% foreground cells **(d)**. User 4 selected 6 cell clusters with 98.6% foreground cells **(e)**. User 5 selected 4 cell clusters with 97.7% foreground cells **(f)**. The label of the clusters selected by using `FlowJo` is in accordance with the colours of the clusters calculated by `flowEMMi`. The mean values and abundances of all cell clusters calculated by `flowEMMi` and `FlowJo` can be found in the additional file `039.csv`.

**Figure 6.**
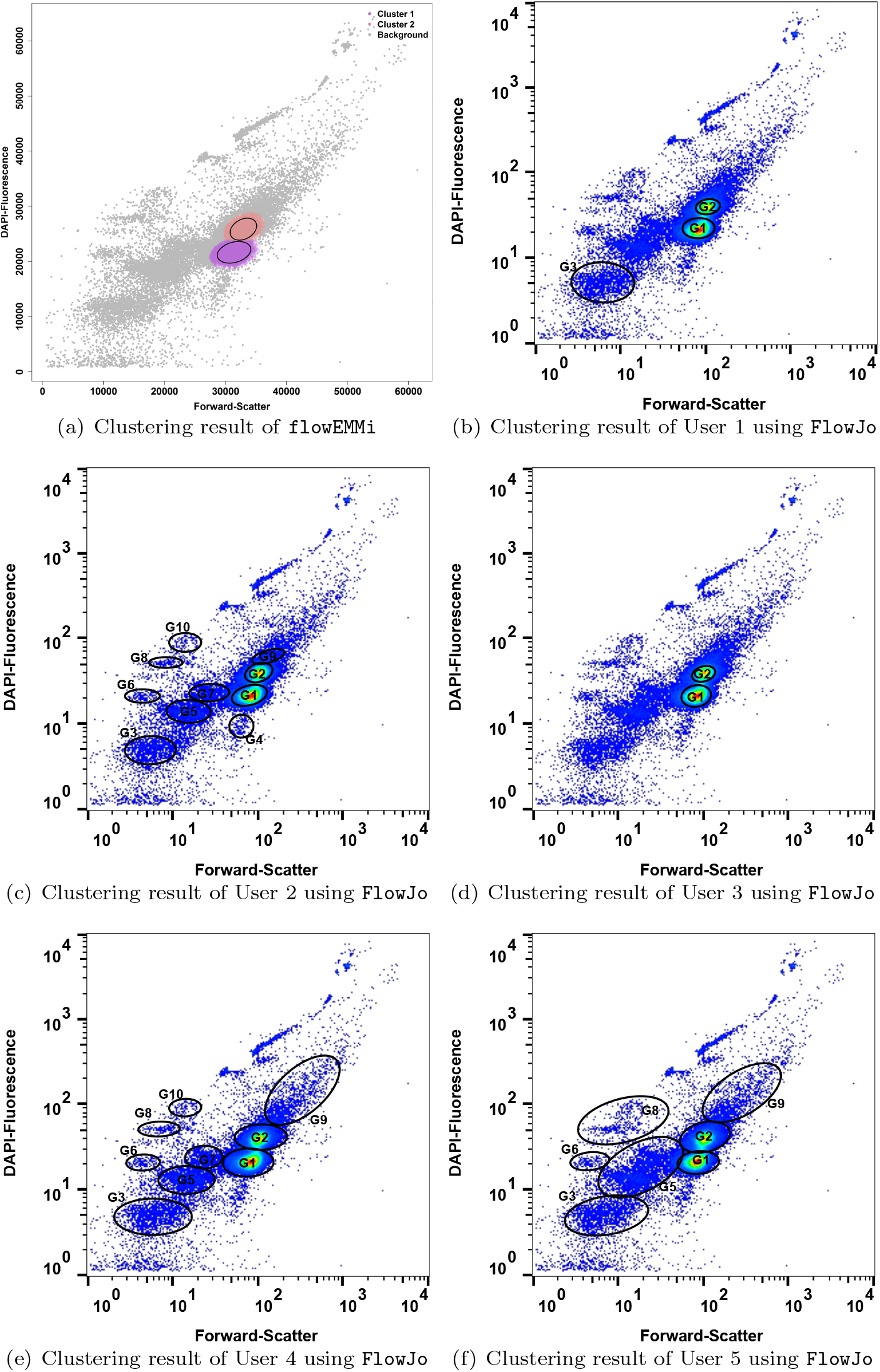
Clustering results for sample InTH_160728_034 using `flowEMMi` with 2 congruent cell clusters and 94.1% foreground cells **(a)** and manual clustering performed by 5 expert users using `FlowJo` **(b-f)**. User 1 selected 3 cell clusters with 88.8% foreground cells **(b)**. User 2 selected 10 cell clusters with 94% foreground cells **(c)**. User 3 selected 2 cell clusters with 88.7% foreground cells **(d)**. User 4 selected 9 cell clusters with 99% foreground cells **(e)**. User 5 selected 7 cell clusters with 100% foreground cells **(f)**. The label of the clusters selected by using FlowJo is in accordance with the colours of the clusters calculated by `flowEMMi`. The mean values and abundances of all cell clusters calculated by `flowEMMi` and `FlowJo` can be found in the additional file `034.csv`.

**Figure 7.**
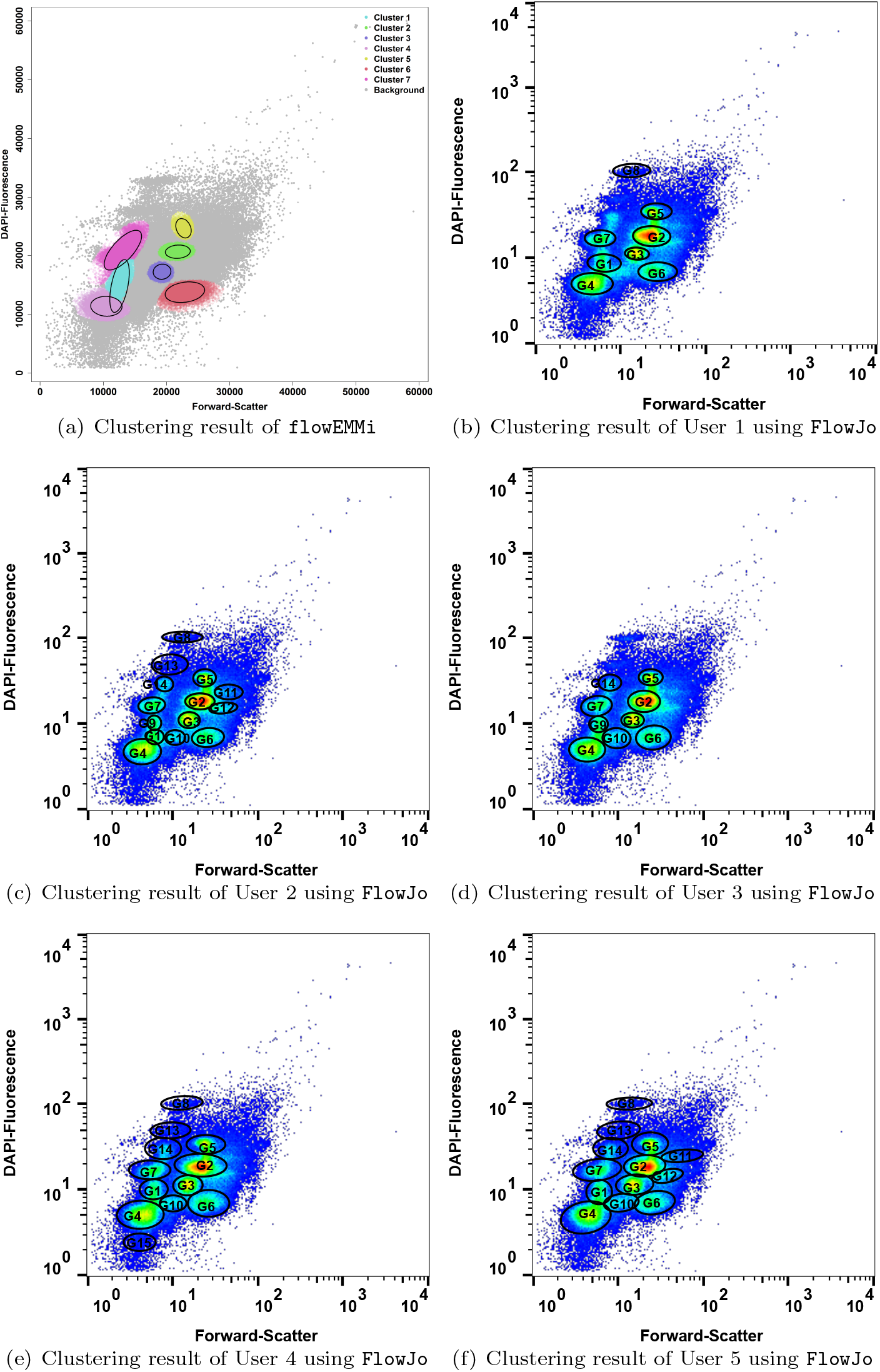
Clustering results for sample InTH_160720_026 using `flowEMMi` with 7 congruent cell clusters and 76.4% foreground cells **(a)** and manual clustering performed by 5 expert users using `FlowJo` **(b-f)**. User 1 selected 8 cell clusters with 76% foreground cells **(b)**. User 2 selected 14 cell clusters with 82.8% foreground cells **(c)**. User 3 selected 9 cell clusters with 79.5% foreground cells **(d)**. User 4 selected 12 cell clusters with 86.9% foreground cells **(e)**. User 5 selected 13 cell clusters with 95.9% foreground cells **(f)**. The label of the clusters selected by using `FlowJo` is in accordance with the colours of the clusters calculated by `flowEMMi`. The mean values and abundances of all cell clusters calculated by `flowEMMi` and `FlowJo` can be found in the additional file `026.csv`.

**Figure 8.**
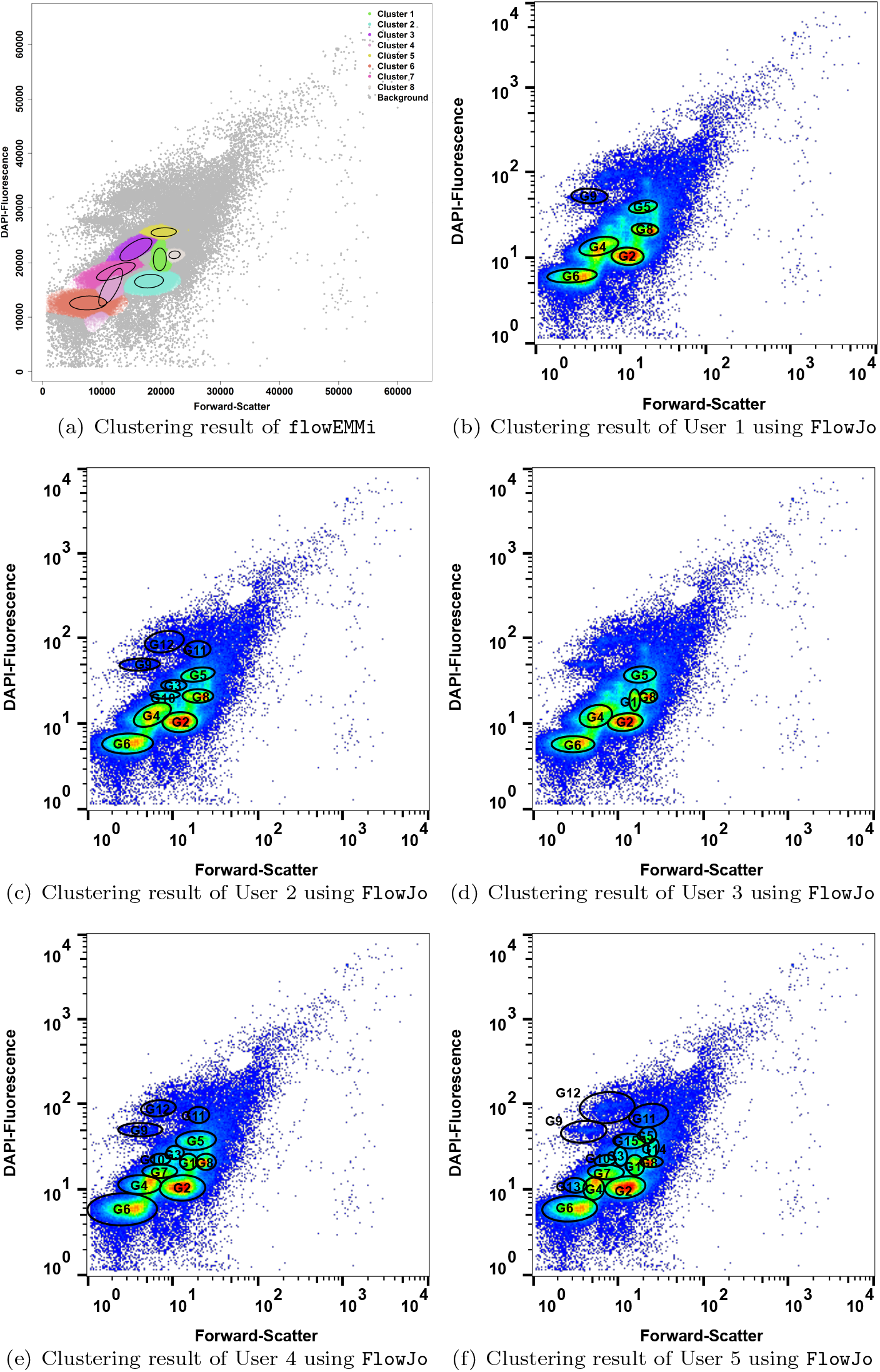
Clustering results for sample InTH_160715_019 using `flowEMMi` with 8 congruent cell clusters and 64.6% foreground cells **(a)** and manual clustering performed by 5 expert users using FlowJo **(b-f)**. User 1 selected 6 cell clusters with 60.1% foreground cells **(b)**. User 2 selected 10 cell clusters with 75.9% foreground cells **(c)**. User 3 selected 6 cell clusters with 67.2% foreground cells **(d)**. User 4 selected 12 cell clusters with 87.7% foreground cells **(e)**. User 5 selected 15 cell clusters with 90.6% foreground cells **(f)**. The label of the clusters selected by using `FlowJo` is in accordance with the colours of the clusters calculated by `flowEMMi`. The mean values and abundances of all cell clusters calculated by `flowEMMi` and `FlowJo` can be found in the additional file `019.csv`.

**Figure 9.**
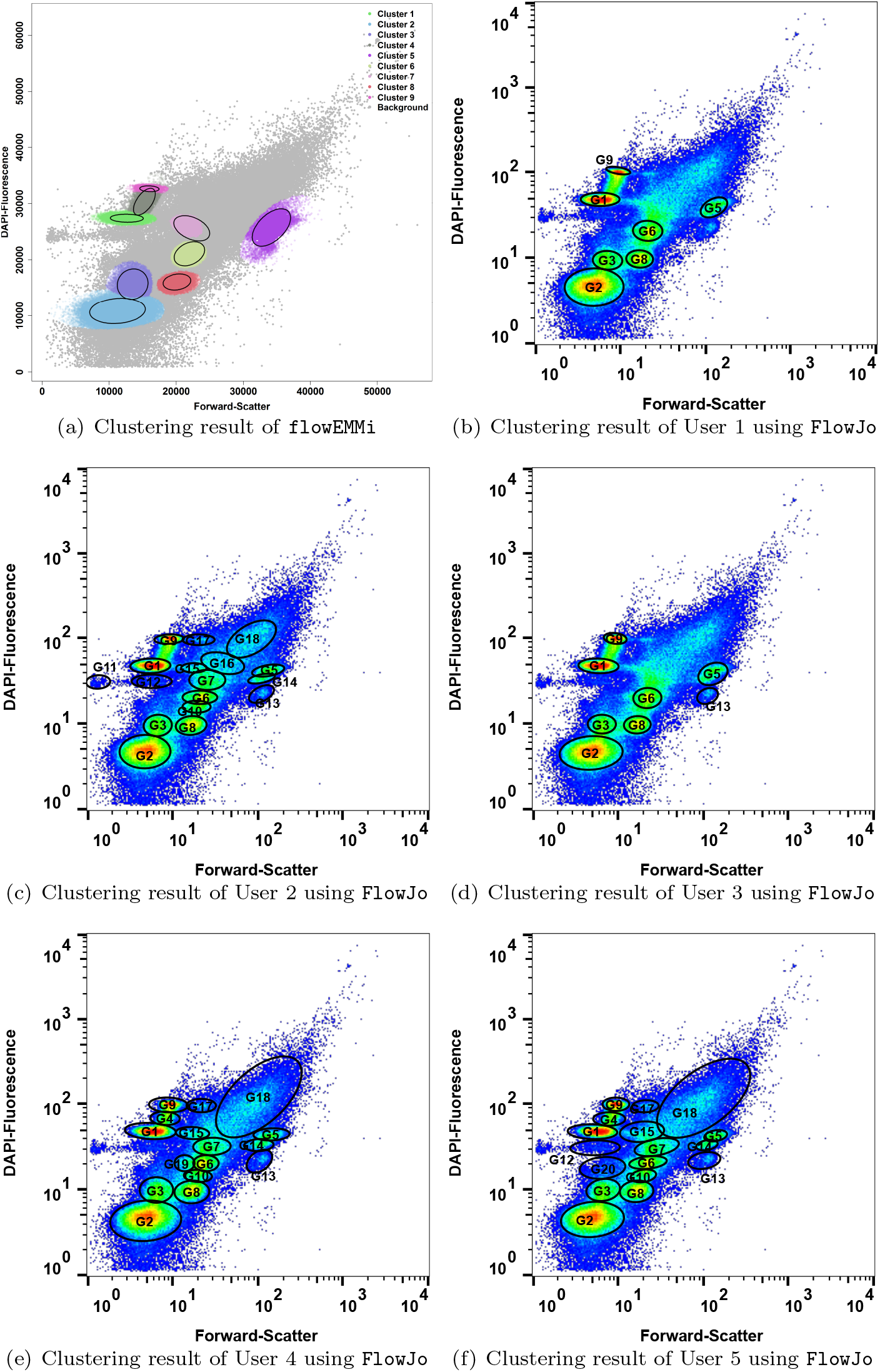
Clustering results for sample InTH_160714_033 using `flowEMMi` with 9 congruent cell clustersand 74.7% foreground cells **(a)** and manual clustering performed by 5 expert users using `FlowJo` **(b-f)**. User 1 selected 7 cell clusters with 61.7% foreground cells **(b)**. User 2 selected 17 cell clusters with 80.1% foreground cells **(c)**. User 3 selected 8 cell clusters with 63.2% foreground cells **(d)**. User 4 selected 16 cell clusters with 92.7% foreground cells **(e)**. User 5 selected 17 cell clusters with 90.2% foreground cells **(f)**. The label of the clusters selected by using FlowJo is in accordance with the colours of the clusters calculated by `flowEMMi`. The mean values and abundances of all cell clusters calculated by `flowEMMi` and `FlowJo` can be found in the additional file `033.csv`.

**Figure 10.**
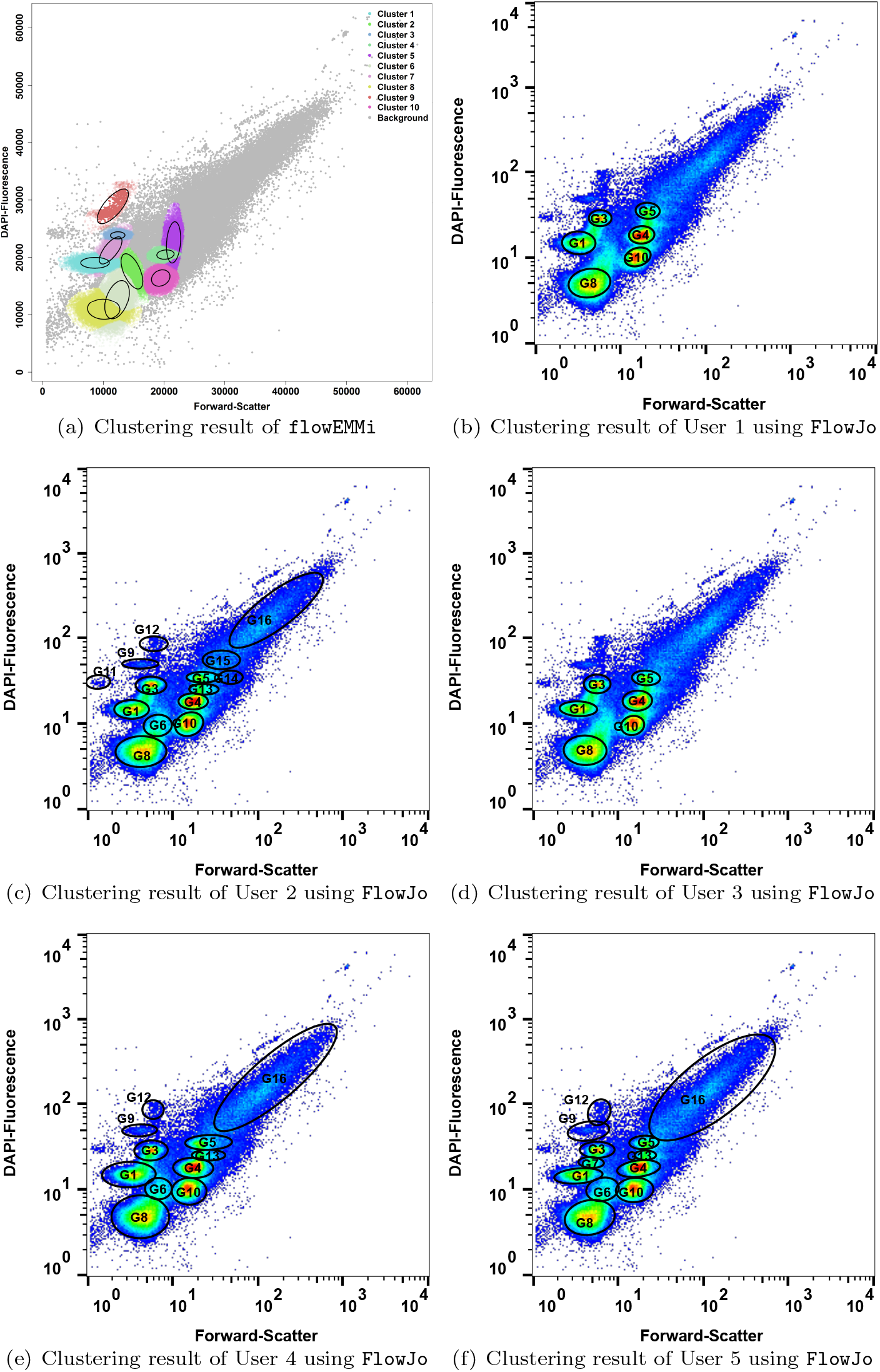
Clustering results for sample InTH_160729_027 using `flowEMMi` with 10 congruent cell clusters and 66.4% foreground cells **(a)** and manual clustering performed by 5 expert users using `FlowJo` **(b-f)**. User 1 selected 6 cell clusters with 69.5% foreground cells **(b)**. User 2 selected 14 cell clusters with 87% foreground cells **(c)**. User 3 selected 6 cell clusters with 69.9% foreground cells **(d)**. User 4 selected 11 cell clusters with 93.7% foreground cells **(e)**. User 5 selected 12 cell clusters with 93% foreground cells **(f)**. The label of the clusters selected by using FlowJo is in accordance with the colours of the clusters calculated by `flowEMMi`. The mean values and abundances of all cell clusters calculated by `flowEMMi` and `FlowJo` can be found in the additional file `027.csv`.

**Figure 11.**
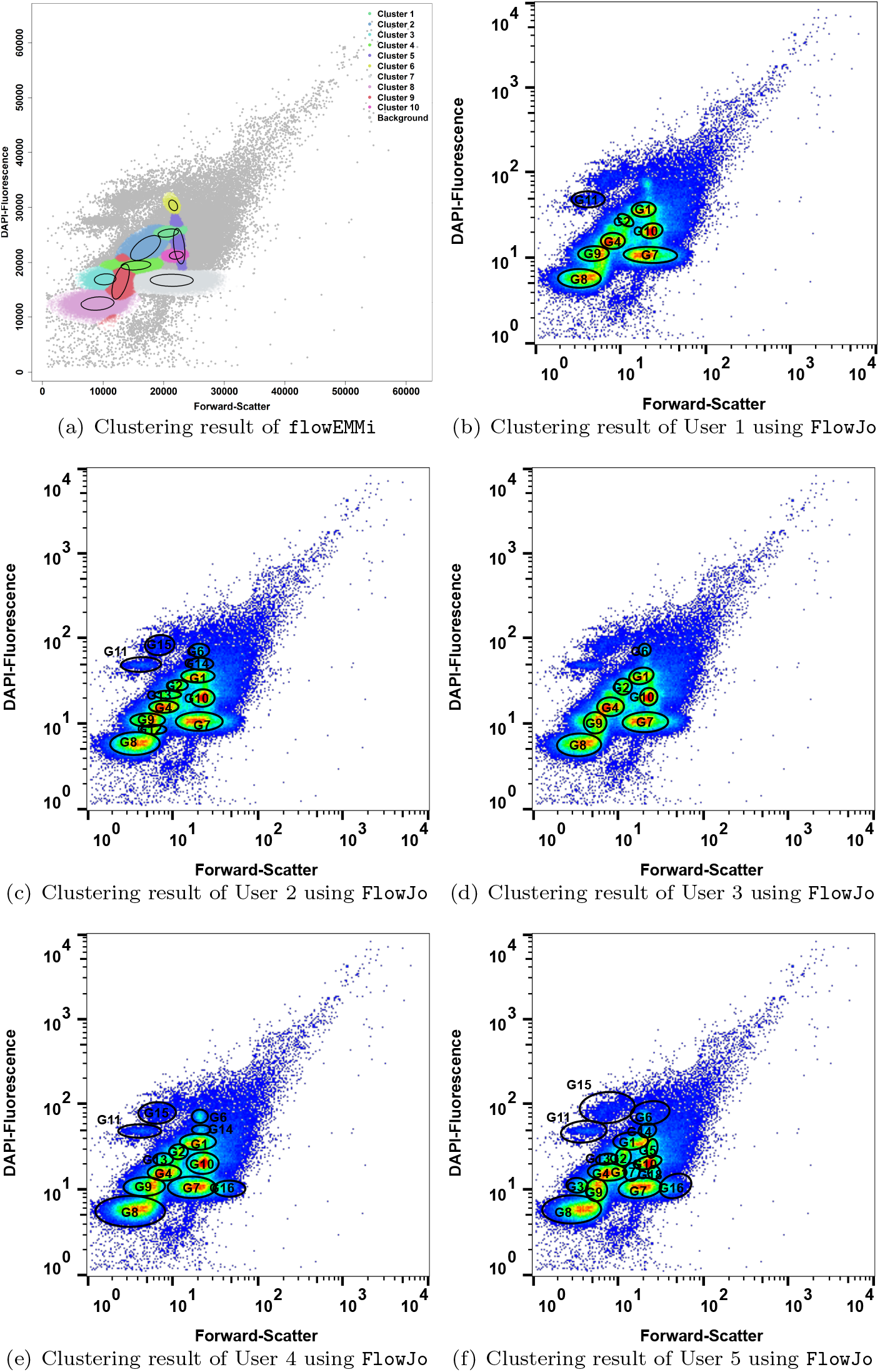
Clustering results for sample InTH_160715_020 using `flowEMMi` with 10 congruent cell clusters and 55.8% foreground cells **(a)** and manual clustering performed by 5 expert users using `FlowJo` **(b-f)**. User 1 selected 8 cell clusters with 64.2% foreground cells **(b)**. User 2 selected 13 cell clusters with 78.2% foreground cells **(c)**. User 3 selected 8 cell clusters with 70.5% foreground cells **(d)**. User 4 selected 13 cell clusters with 86.8% foreground cells **(e)**. User 5 selected 17 cell clusters with 91.3% foreground cells **(f)**. The label of the clusters selected by using `FlowJo` is in accordance with the colours of the clusters calculated by `flowEMMi`. The mean values and abundances of all cell clusters calculated by `flowEMMi` and `FlowJo` can be found in the additional file `020.csv`.

**Figure 12.**
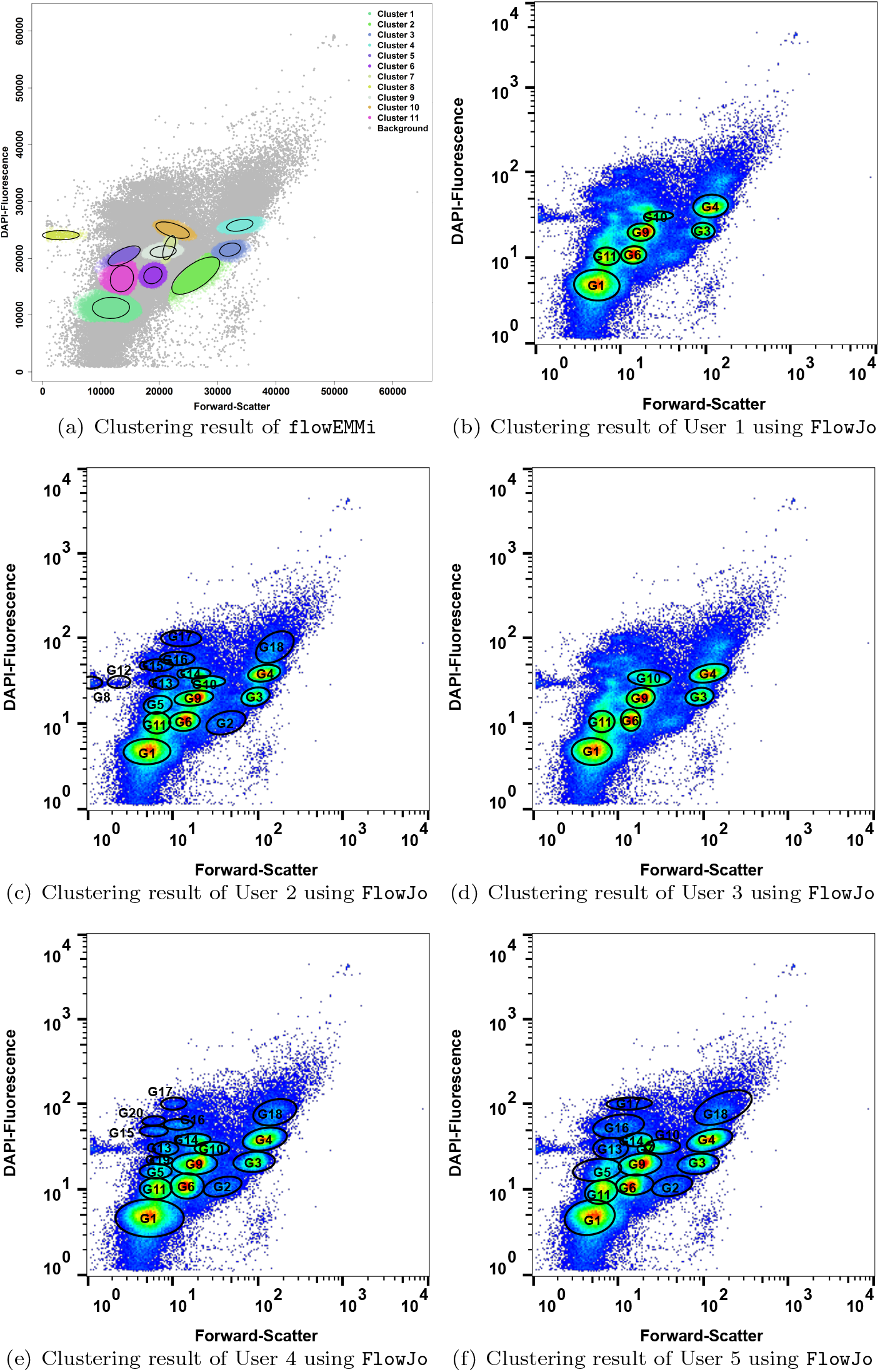
Clustering results for sample InTH_160720_035 using `flowEMMi` with 11 congruent cell clusters and 72.6% foreground cells **(a)** and manual clustering performed by 5 expert users using `FlowJo` **(b-f)**. User 1 selected 7 cell clusters with 69.5% foreground cells **(b)**. User 2 selected 17 cell clusters with 81.7% foreground cells **(c)**. User 3 selected 7 cell clusters with 71.5% foreground cells **(d)**. User 4 selected 17 cell clusters with 88.5% foreground cells **(e)**. User 5 selected 15 cell clusters with 92% foreground cells **(f)**. The label of the clusters selected by using `FlowJo` is in accordance with the colours of the clusters calculated by `flowEMMi`. The mean values and abundances of all cell clusters calculated by `flowEMMi` and `FlowJo` can be found in the additional file `035.csv`.

**Figure 13.**
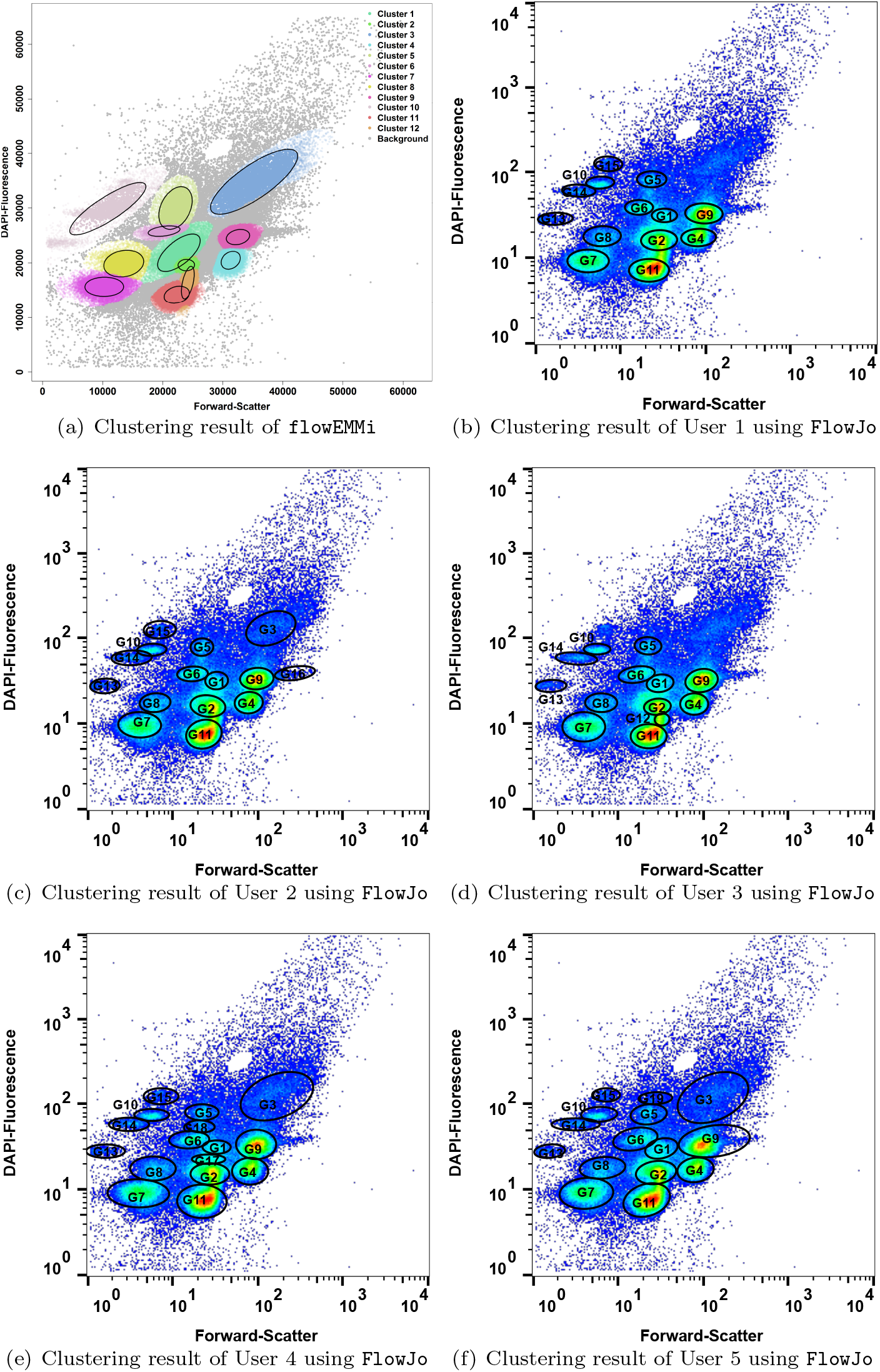
Clustering results for sample InTH_160712_025 using `flowEMMi` with 12 congruent cell clusters and 71.6% foreground cells **(a)** and manual clustering performed by 5 expert users using `FlowJo` **(b-f)**. User 1 selected 13 cell clusters with 76.5% foreground cells **(b)**. User 2 selected 15 cell clusters with 82.1% foreground cells **(c)**. User 3 selected 13 cell clusters with 79.1% foreground cells **(d)**. User 4 selected 16 cell clusters with 90.7% foreground cells **(e)**. User 5 selected 15 cell clusters with 91.6% foreground cells **(f)**. The label of the clusters selected by using `FlowJo` is in accordance with the colours of the clusters calculated by `flowEMMi`. The mean values and abundances of all cell clusters calculated by `flowEMMi` and `FlowJo` can be found in the additional file `025.csv`.

**Figure 14.**
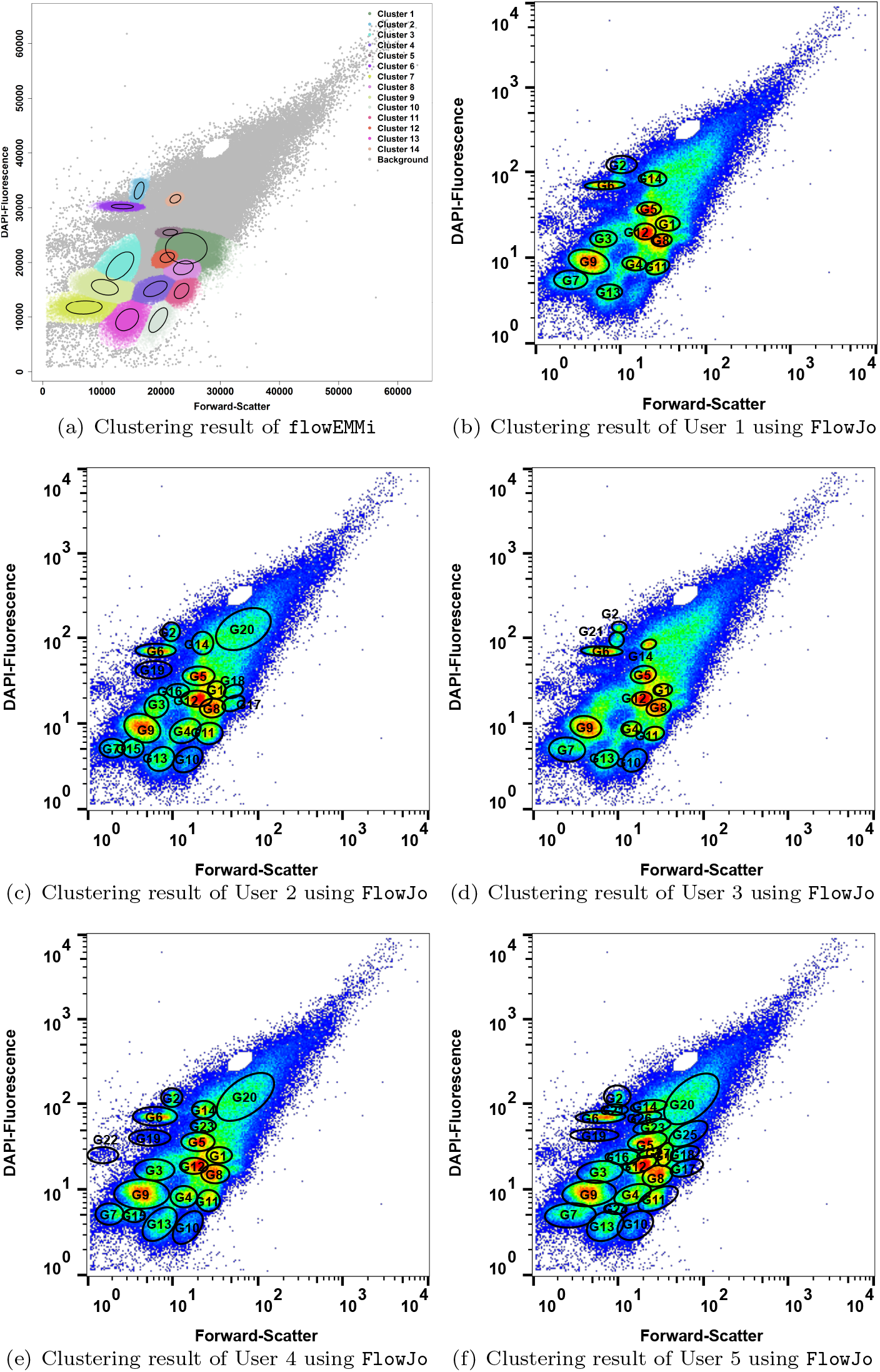
Clustering results for sample InTH_160713_012 using `flowEMMi` with 14 congruent cell clusters and 49.5% foreground cells **(a)** and manual clustering performed by 5 expert users using `FlowJo` **(b-f)**. User 1 selected 13 cell clusters with 49.3% foreground cells **(b)**. User 2 selected 20 cell clusters with 75.4% foreground cells **(c)**. User 3 selected 14 cell clusters with 47.3% foreground cells **(d)**. User 4 selected 19 cell clusters with 66.3% foreground cells **(e)**. User 5 selected 25 cell clusters with 92% foreground cells **(f)**. The label of the clusters selected by using `FlowJo` is in accordance with the colours of the clusters calculated by `flowEMMi`. The mean values and abundances of all cell clusters calculated by `flowEMMi` and `FlowJo` can be found in the additional file `012.csv`.

**Figure 15.**
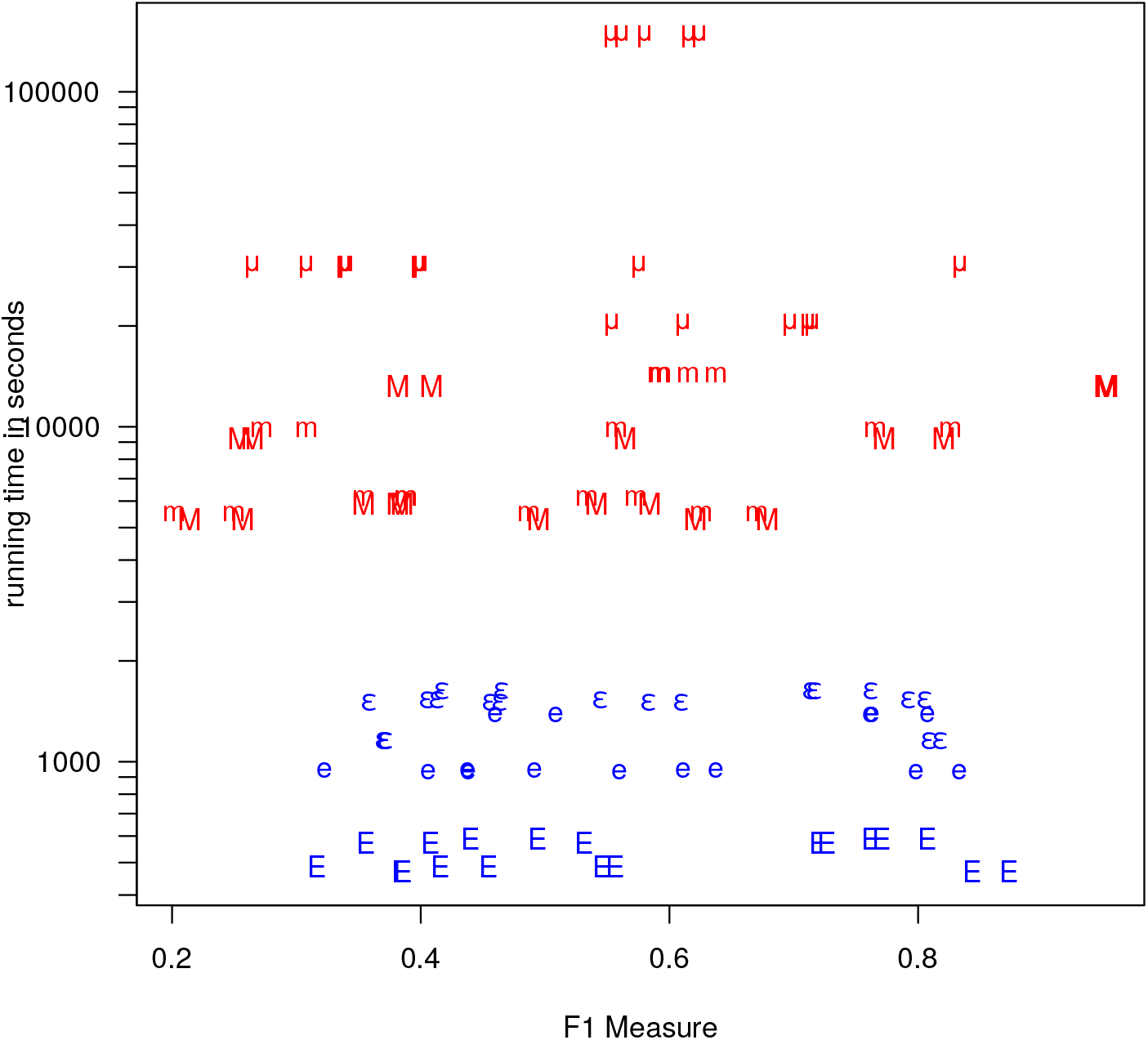
Comparison of running time (note the logarithmic scaling) vs F_1_ score for `flowEMMi` (characters *ϵ*, `e`, `E` in blue) and `flowMerge` (characters *μ*, `m`, `M` in red). `flowEMMi` yields, on average over all runs shown above ≈ 10.7% better F_1_ scores (`flowEMMi` mean: 0.57, sd: 0.17; `flowMerge` mean: 0.53, sd: 0.20), at very different time scales (`flowEMMi` mean: 1012, sd: 427; `flowMerge` mean: 24 665, sd: 38 019). `flowEMMi` performs extremely well in a time constrained regime at early Expectation-Minimization cutoff (using on the cutoff at < 1 instead of cutoff < 0.01 or < 10^−5^) with F_1_ score mean: 0.56, sd:0.18, and a running time in seconds of mean: 528, sd: 53. While `flowMerge` has slightly worse F_1_ score characteristics (mean: 0.54, sd: 0.24), with running times a lot higher (mean: 8 391, sd: 3 239). Since both algorithms are parallelized, actual wall-clock times are lower by a factor of 2–3 on a 4-core machine. Given running times are total core seconds used.

**Figure 16.**
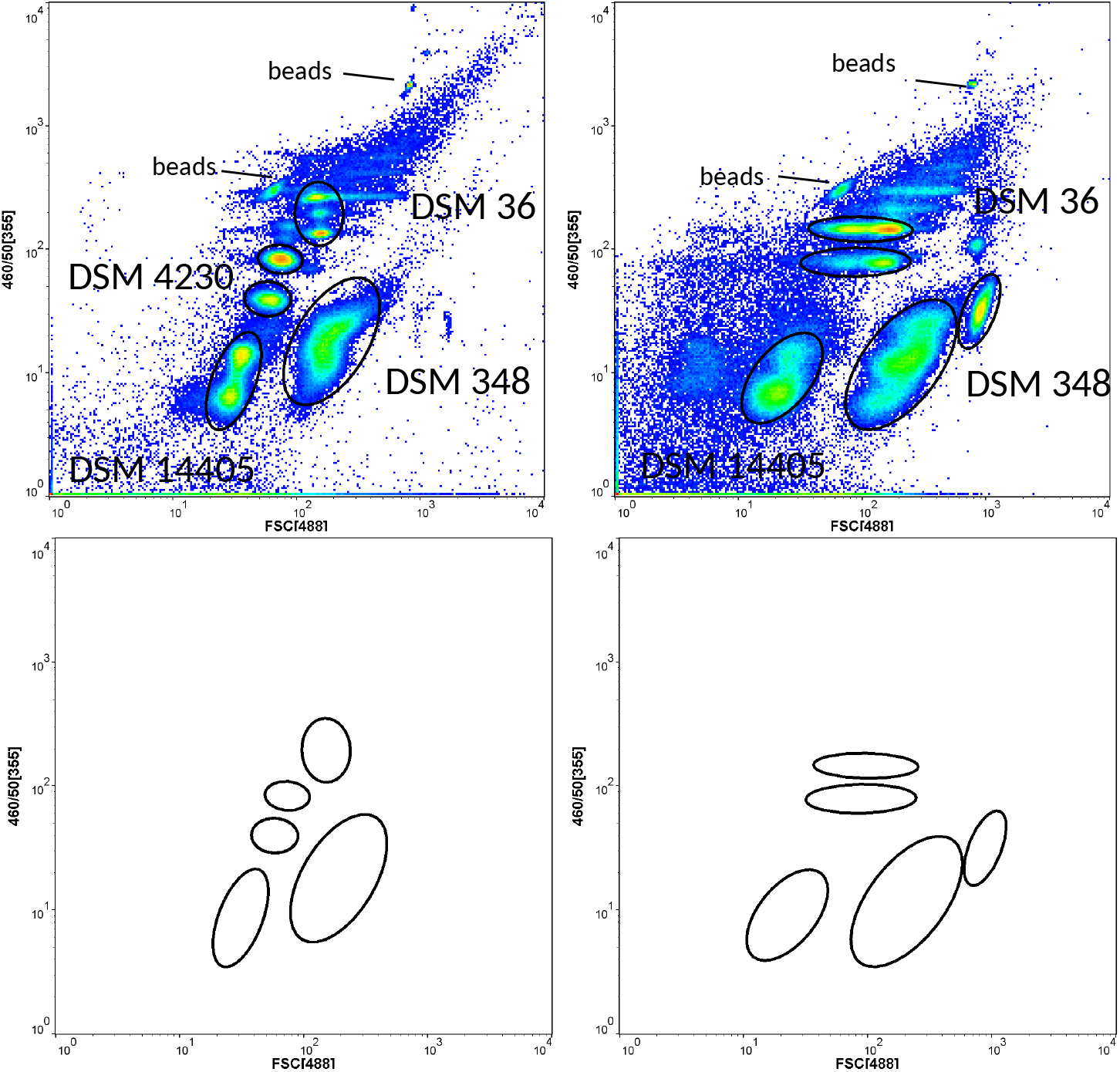
Flow cytometric measurement of microbial cytometric mock communities. **Left:** strains *Stenotrophomonas rhizophila* DSM 14405, *Escherichia coli* DSM 4230, *Kocuria rhizophila* DSM 348, and *Paenibacillus polymyxa* DSM 36 were grown in liquid culture, respectively. **Right:** strains *Stenotrophomonas rhizophila* DSM 14405, *Kocuria rhizophila* DSM 348, and *Paenibacillus polymyxa* DSM 36 were grown on plate. The beads were introduced for instrumental alignment of the flow cytometer. **Below:** manually set gate templates for the liquid (left) and plate (right) microbial cytometric mock communities.

**Figure 17.**
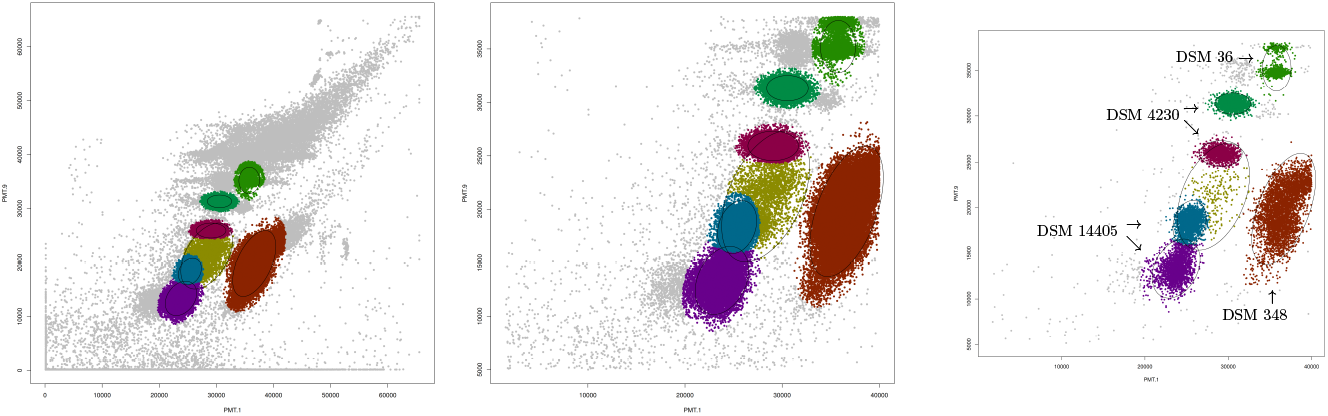
Automated gating by `flowEMMi` revealed the highest abundant subpopulations of the four strains *Stenotrophomonas rhizophila* DSM 14405, *Escherichia coli* DSM 4230, *Kocuria rhizophila* DSM 348, and *Paenibacillus polymyxa* DSM 36 grown in liquid culture. From left to right: full data set, including noise, rectangular cutout without corner noise, gating on subset of data. Automatic gating by `flowEMMi` yields an F_1_ value of 0.85.

**Figure 18.**
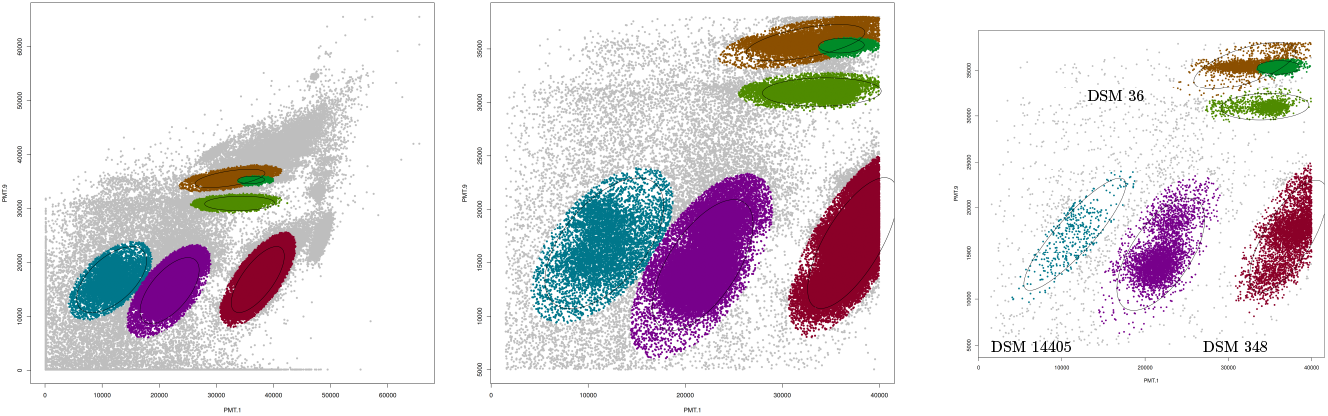
Automated gating by `flowEMMi` revealed the highest abundant subpopulations of the three strains *Stenotrophomonas rhizophila* DSM 14405, *Kocuria rhizophila* DSM 348, and *Paenibacillus polymyxa* DSM 36 grown on plate. From left to right: full data set, including noise, rectangular cutout without corner noise, gating on subset of data. Automatic gating by `flowEMMi` yields an F_1_ value of 0.81.

## References

1. Hammes, F., Berney, M., Wang, Y., Vital, M., Köster, O., Egli, T.: Flow-cytometric total bacterial cell counts as a descriptive microbiological parameter for drinking water treatment processes. Water Research 42(1-2), 269–277 (2008)

2. Lautenschlager, K., Boon, N., Wang, Y., Egli, T., Hammes, F.: Overnight stagnation of drinking water in household taps induces microbial growth and changes in community composition. Water Research 44(17), 4868–4877 (2010)

3. Lautenschlager, K., Hwang, C., Ling, F., Liu, W.-T., Boon, N., Köster, O., Egli, T., Hammes, F.: Abundance and composition of indigenous bacterial communities in a multi-step biofiltration-based drinking water treatment plant. Water Research 62, 40–52 (2014)

4. Günther, S., Faust, K., Schumann, J., Harms, H., Raes, J., Müller, S.: Species-sorting and mass-transfer paradigms control managed natural metacommunities. Environmental Microbiology 18(12), 4862–4877 (2016)

5. Lambrecht, J., Cichocki, N., Hübschmann, T., Koch, C., Harms, H., Müller, S.: Flow cytometric quantification, sorting and sequencing of methanogenic archaea based on F 420 autofluorescence. Microbial Cell Factories 16(1), 180 (2017)

6. Props, R., Monsieurs, P., Mysara, M., Clement, L., Boon, N.: Measuring the biodiversity of microbial communities by flow cytometry. Methods in Ecology and Evolution 7(11), 1376–1385 (2016)

7. Liu, Z., Cichocki, N., Bonk, F., Günther, S., Schattenberg, F., Harms, H., Centler, F., Müller, S.: Ecological stability properties of microbial communities assessed by flow cytometry. mSphere 3(1), 00564–17 (2018)

8. Liu, Z., Cichocki, N., Hübschmann, T., Süring, C., Ofiţeru, I.D., Sloan, W.T., Grimm, V., Müller, S.: Neutral mechanisms and niche differentiation in steady-state insular microbial communities revealed by single cell analysis. Environmental microbiology (2019)

9. Zimmermann, J., Hübschmann, T., Schattenberg, F., Schumann, J., Durek, P., Riedel, R., Friedrich, M., Glauben, R., Siegmund, B., Radbruch, A., Müller, S., Chang, H.-D.: High-resolution microbiota flow cytometry reveals dynamic colitis-associated changes in fecal bacterial composition. European Journal of Immunology 46(5), 1300–1303 (2016)

10. van Gelder, S., Röhrig, N., Schattenberg, F., Cichocki, N., Schumann, J., Schmalz, G., Haak, R., Ziebolz, D., Müller, S.: A cytometric approach to follow variation and dynamics of the salivary microbiota. Methods 134-135, 67–79 (2018)

11. Koch, C., Müller, S.: Personalized microbiome dynamics–cytometric fingerprints for routine diagnostics. Molecular aspects of medicine 59, 123–134 (2018)

12. Buysschaert, B., Kerckhof, F.-M., Vandamme, P., De Baets, B., Boon, N.: Flow cytometric fingerprinting for microbial strain discrimination and physiological characterization. Cytometry Part A 93(2), 201–212 (2018)

13. Rubbens, P., Props, R., Boon, N., Waegeman, W.: Flow cytometric single-cell identification of populations in synthetic bacterial communities. PloS one 12(1), 0169754 (2017)

14. Schumann, J., Koch, C., Fetzer, I., Müller, S.: flowCHIC: Analyze Flow Cytometric Data Using Histogram Information. (2015). R package version 1.6.0

15. Koch, C., Fetzer, I., Harms, H., Müller, S.: CHIC - an automated approach for the detection of dynamic variations in complex microbial communities. Cytometry Part A 83(6), 561–567 (2013)

16. Schumann, J., Koch, C., Günther, S., Fetzer, I., Müller, S.: flowCyBar: Analyze Flow Cytometric Data Using Gate Information. (2015). R package version 1.8.0

17. Koch, C., Fetzer, I., Schmidt, T., Harms, H., Müller, S.: Monitoring functions in managed microbial systems by cytometric bar coding. Environmental science & technology 47(3), 1753–1760 (2013)

18. De Novo Software: FCS Express. https://www.denovosoftware.com/

19. Sysmex Partec GmbH: FloMax. https://www.sysmex-partec.com/

20. Dako Colorado Inc.: Summit. https://www.med.unc.edu/flowcytometry/instrumentation-2/data-analysis/download-summit

21. Mehta, T., Bose, B.: FlowPy. (2010). http://flowpy.wikidot.com

22. Aghaeepour, N., Finak, G., Hoos, H., Mosmann, T.R., Brinkman, R., Gottardo, R., Scheuermann, R.H., FlowCAP Consortium, DREAM Consortium: Critical assessment of automated flow cytometry data analysis techniques. Nature Methods 10(3), 228–238 (2013)

23. Günther, S., Müller, S.: Facilitated gate setting by sequential dot plot scanning. Cytometry Part A 87(7), 661–664 (2015)

24. Lo, K., Hahne, F., Brinkman, R.R., Gottardo, R.: flowClust: a Bioconductor package for automated gating of flow cytometry data. BMC Bioinformatics 10(1), 145 (2009)

25. FlowJo, LLC: FlowJo. https://www.flowjo.com/

26. Holyst, H., Rogers, W.: flowFP: Fingerprinting for Flow Cytometry. (2009). R package version 1.30.0

27. Roederer, M., Moore, W., Treister, A., Hardy, R.R., Herzenberg, L.A.: Probability binning comparison: A metric for quantitating multivariate distribution differences. Cytometry Part A 45(1), 47–55 (2001)

28. Zare, H., Shooshtari, P., Gupta, A., Brinkman, R.R.: Data reduction for spectral clustering to analyze high throughput flow cytometry data. BMC Bioinformatics 11(1), 403 (2010)

29. Kanungo, T., Mount, D.M., Netanyahu, N.S., Piatko, C.D., Silverman, R., Wu, A.Y.: An efficient k-means clustering algorithm: Analysis and implementation. IEEE transactions on pattern analysis and machine intelligence 24(7), 881–892 (2002)

30. Malek, M., Taghiyar, M.J., Chong, L., Finak, G., Gottardo, R., Brinkman, R.R.: flowDensity: reproducing manual gating of flow cytometry data by automated density-based cell population identification. Bioinformatics 31(4), 606–607 (2014)

31. Aghaeepour, N., Nikolic, R., Hoos, H.H., Brinkman, R.R.: Rapid cell population identification in flow cytometry data. Cytometry Part A 79(1), 6–13 (2011)

32. Finak, G., Bashashati, A., Brinkman, R., Gottardo, R.: Merging mixture components for cell population identification in flow cytometry. Advances in bioinformatics 2009 (2009)

33. Pyne, S., Hu, X., Wang, K., Rossin, E., Lin, T.-I., Maier, L.M., Baecher-Allan, C., McLachlan, G.J., Tamayo, P., Hafler, D.A., De Jager, P.L., Mesirov, J.P.: Automated high-dimensional flow cytometric data analysis. Proceedings of the National Academy of Sciences 106(21), 8519–8524 (2009)

34. Brinkman, R.R., Gasparetto, M., Lee, S.-J.J., Ribickas, A.J., Perkins, J., Janssen, W., Smiley, R., Smith, C.: High-content flow cytometry and temporal data analysis for defining a cellular signature of graft-versus-host disease. Biology of Blood and Marrow Transplantation 13(6), 691–700 (2007)

35. Amalfitano, S., Fazi, S., Ejarque, E., Freixa, A., Romani, A.M., Butturini, A.: Deconvolution model to resolve cytometric microbial community patterns in flowing waters. Cytometry Part A 93(2), 194–200 (2018)

36. Koch, C., Günther, S., Desta, A.F., Hübschmann, T., Müller, S.: Cytometric fingerprinting for analyzing microbial intracommunity structure variation and identifying subcommunity function. Nature Protocols 8(1), 190–202 (2013)

37. Shapiro, H.M.: Practical Flow Cytometry. John Wiley & Sons, Hoboken (New Jersey) (2005)

38. Müller, S., Nebe-von-Caron, G.: Functional single-cell analyses: flow cytometry and cell sorting of microbial populations and communities. FEMS Microbiology Reviews 34(4), 554–587 (2010)

39. Baudry, J.-P., Celeux, G.: EM for mixtures. Statistics and computing 25(4), 713–726 (2015)

40. Yu, J., Qin, S.J.: Multimode process monitoring with bayesian inference-based finite Gaussian mixture models. AIChE Journal 54(7), 1811–1829 (2008)

41. Dempster, A.P., Laird, N.M., Rubin, D.B.: Maximum likelihood from incomplete data via the EM algorithm. Journal of the royal statistical society. Series B (methodological), 1–38 (1977)

42. Wu, C.J.: On the convergence properties of the EM algorithm. The Annals of statistics, 95–103 (1983)

43. Connor, R.J., Mosimann, J.E.: Concepts of independence for proportions with a generalization of the Dirichlet distribution. Journal of the American Statistical Association 64(325), 194–206 (1969)

44. Eddelbuettel, D., François, R.: Rcpp: Seamless R and C++ integration. Journal of Statistical Software 40(8), 1–18 (2011). doi:10.18637/jss.v040.i08

45. Bates, D., Eddelbuettel, D.: Fast and elegant numerical linear algebra using the RcppEigen package. Journal of Statistical Software 52(5), 1–24 (2013)

46. Ellis, B., Haaland, P., Hahne, F., Le Meur, N., Gopalakrishnan, N., Spidlen, J., Jiang, M.: flowCore: Basic Structures for Flow Cytometry Data. (2016). R package version 1.38.2

47. Ellis, B., Gentleman, R., Hahne, F., Le Meur, N., Sarkar, D., Jiang, M.: flowViz: Visualization for Flow Cytometry. (2016). R package version 1.36.2

48. Wickham, H.: ggplot2: Elegant Graphics for Data Analysis. Springer, New York (2009). http://ggplot2.org

49. Ammar, R.: randomcoloR: Generate Attractive Random Colors. (2016). R package version 1.0.0. https://CRAN.R-project.org/package=randomcoloR

50. Benaglia, T., Chauveau, D., Hunter, D.R., Young, D.: mixtools: An R package for analyzing finite mixture models. Journal of Statistical Software 32(6), 1–29 (2009)

51. Warnes, G.R., Bolker, B., Lumley, T.: gtools: Various R Programming Tools. (2015). R package version 3.5.0. https://CRAN.R-project.org/package=gtools

52. Schwarz, G.: Estimating the dimension of a model. The annals of statistics 6(2), 461–464 (1978)

53. Wit, E., Heuvel, E.v.d., Romeijn, J.-W.: ‘all models are wrong…’: an introduction to model uncertainty. Statistica Neerlandica 66(3), 217–236 (2012)

54. Neal, R.M., Hinton, G.E.: A view of the EM algorithm that justifies incremental, sparse, and other variants. In: Learning in Graphical Models, pp. 355–368. Springer, New York (1998)

55. Neyman, J.: Outline of a theory of statistical estimation based on the classical theory of probability. Phil. Trans. R. Soc. Lond. A 236(767), 333–380 (1937)

56. Zar, J.H.: Biostatistical analysis. 2nd. Prentice Hall USA 54, 55 (1984)

57. Koch, C., Müller, S.: Personalized microbiome dynamics - Cytometric fingerprints for routine diagnostics. Molecular Aspects of Medicine 59, 123–134 (2018)

